# Paclitaxel-induced mitotic arrest results in a convergence of apoptotic dependencies that can be safely exploited by BCL-X_L_ degradation to overcome cancer chemoresistance

**DOI:** 10.1101/2025.06.24.661170

**Authors:** Xingping Qin, Adam Presser, Lissah Johnson, Yusuke Matoba, Brendan S. Shay, Weiyi Xu, Jonathan Choiniere, Cameron Fraser, Filip Garbicz, Johan Spetz, Stacey Yu, Mary H. C. Florido, Francesca Picucci, Yang Yang, Ronny Drapkin, Ruben Carrasco, Sarah J. Hill, Joyce Liu, Ursula Matulonis, Joan Brugge, Bo R. Rueda, Daohong Zhou, Elizabeth H. Stover, Kristopher A. Sarosiek

## Abstract

Paclitaxel and other microtubule-targeting agents are cornerstone therapies for diverse cancers, including lung, breast, cervical, pancreatic, and ovarian malignancies. Paclitaxel induces tumor cell apoptosis during mitosis by disrupting microtubule dynamics required for chromosome segregation. However, despite initial responsiveness, many tumors develop resistance, limiting therapeutic durability. Here, we used high-grade serous ovarian carcinoma (HGSOC), the most common and lethal subtype of ovarian cancer, as a model to dissect the mechanisms underlying this resistance. We find that paclitaxel-induced mitotic arrest triggers degradation of the pro-survival protein MCL-1 and upregulation of BCL-XL, followed by inactivating phosphorylation of BCL-XL at Ser62 to promote apoptosis. In resistant cells, this MCL-1 downregulation is insufficient to commit cells to apoptosis but instead results in a transient convergence of apoptotic dependencies by forcing BCL-X_L_ to sequester the pro-apoptotic proteins BIM, BAX, and BAK. During this state, BCL-XL inhibition induces synergistic apoptosis, even in chemoresistant cells. Surprisingly, we also discover that loss of substrate attachment recapitulates this apoptotic convergence both in vitro and in vivo, with HGSOC cells growing in metastasis-promoting malignant ascites displaying heightened apoptotic priming and dependence on BCL-XL relative to solid tumors. In HGSOC xenografts, targeted degradation of BCL-XL using the platelet-sparing proteolysis-targeting chimera (PROTAC) DT2216 matches the efficacy of paclitaxel monotherapy while avoiding the chronic thrombocytopenia induced by BCL-XL inhibitors such as navitoclax (ABT-263). Strikingly, combination therapy leveraging the synergy between paclitaxel and DT2216 leads to complete eradication of HGSOC cell line and patient-derived xenografts. Moreover, DT2216 treatment blunts the rapid apoptotic adaptation caused by other BCL-X_L_ inhibitors, indicating that targeted degradation of pro-survival proteins may yield more durable responses than inhibition alone. These findings uncover a mechanistic framework for safely exploiting the apoptotic dependency convergence caused by mitotic arrest and substrate detachment and support the clinical development of BCL-XL–targeting PROTACs to overcome chemoresistance in ovarian cancer and other solid tumors.

## Introduction

Ovarian cancers (OvCa) of epithelial origin comprise a heterogeneous group of malignancies that are usually diagnosed in patients as an advanced disease^1^. The most common and clinically intractable of these are high-grade serous ovarian carcinomas (HGSOC), which exhibit an alarmingly low 5-year overall survival (OS) rate of 35-48%^2,3^. This OS rate has not meaningfully improved in the past 20 years due to a lack of clinically actionable targets, as well as the profound genomic instability that is a hallmark of this disease. This instability is driven by near-ubiquitous mutations in *TP53* as well as deficiencies in homologous recombination and other DNA repair pathways in a significant percentage of HGSOC cases^2,4^ – these features contribute to the development of chemoresistance if the tumor is not eradicated upon initial treatment. Treatment with standard therapy, which includes carboplatin and paclitaxel, most commonly results in a period of cancer remission of variable length, typically 6 months to several years^5–7^. In 60-80% of patients, this period is followed by tumor recurrence, with each subsequent recurrent tumor exhibiting increased resistance to chemotherapy and almost no possibility of cure^6,7^. These high rates of relapse and mortality demonstrate the urgent need to develop treatments that can more thoroughly eradicate HGSOC cells to prevent the development of therapy resistance and increase OS rates. Notably, carboplatin and paclitaxel target ubiquitous cell features, including DNA and microtubules, respectively, and typically kill OvCa cells via the apoptosis pathway^8–10^

Apoptosis is a regulated form of cell death that is essential for normal human development as well as for the removal of damaged or superfluous cells^11–14^. The mitochondrial apoptosis pathway is triggered when a pro-apoptotic protein (BAX or BAK) is activated by a BH3-only protein (BIM or BID) to trigger mitochondrial outer membrane permeabilization (MOMP) and consequent release of cytochrome *c* from mitochondria to activate caspases^14–16^. Pro-survival proteins in this family (predominantly BCL-2, BCL-X_L_, and MCL-1) can bind and block the activity of either the pro-apoptotic, pore-forming proteins (BAX or BAK) or pro-apoptotic BH3-only activators (BIM, BID, PUMA). For apoptosis to occur, the pro-survival proteins must be overwhelmed and BAX/BAK activated.

Importantly, we have previously reported that the state of the apoptosis pathway in HGSOC is an important determinant of chemotherapy response: tumors that are primed for apoptosis respond much more favorably in the clinic^17–19^. Furthermore, we and others have shown that upregulation of pro-survival BCL-X_L_ or MCL-1^20–23^ or loss of pro-apoptotic BAX and BAK^18,24,25^ are major drivers of therapy resistance in HGSOC. These findings, along with those of others^23,26,27^ suggest that patient outcomes can be meaningfully improved by targeting the apoptosis pathway to enhance initial chemotherapy responses. BH3 mimetics are recently-developed agents that selectively inhibit certain pro-survival proteins including BCL-2 (e.g., ABT-199), BCL-X_L_ (e.g., A1331852) or MCL-1 (e.g., S63845) and have been successful in the clinic for targeting apoptotic dependencies in hematologic malignancies^28^. However, although the BCL-2 inhibitor ABT-199 (venetoclax) is extremely well tolerated and FDA approved as a treatment for chronic lymphocytic leukemia and acute myeloid leukemia, inhibitors targeting other pro-survival proteins do cause some on-target toxicities. For example, existing small molecule inhibitors of BCL-X_L_ are known to be toxic to platelets due to their dependence on BCL-X_L_ for survival. This causes dose-limiting thrombocytopenia and increases risk of bleeding, which has impeded the clinical development of these agents. To overcome this, DT2216, a BCL-X_L_ inhibiting proteolysis-targeting chimera (PROTAC) has been developed that degrades BCL-X_L_ by targeting it to the Von Hippel-Lindau (VHL) E3 ligase for degradation instead of simply inhibiting it – this is an important modification as platelets express minimal levels of VHL and therefore don’t degrade BCL-X_L_ in a manner similar to most nucleated cells. This prevents the loss of platelets and increases the safety of this novel agent^29,30^, which is currently being evaluated in phase I/II clinical trials (NCT06620302).

Although there have been promising recent advancements in clinical deployment of these BH3 mimetics, it is unknown what apoptotic vulnerabilities exist in HGSOC and how they may be effectively and safely targeted with single agent inhibitors or in combinations. Herein, we report that BCL-X_L_ dependence is consistently detected in a set of HGSOC cell lines, primary tumors and PDX models. Strikingly, treatment of HGSOC primary tumors and PDX models with the standard of care agent paclitaxel primes HGSOC for apoptosis and enhances sensitivity to BCL-X_L_ inhibition, leading to eradication of tumors in vivo while avoiding toxicities.

## Results

### Epithelial ovarian cancer cell lines are primed for apoptosis and dependent on BCL-X_L_ for survival

We first sought to measure apoptotic priming and uncover potential dependencies on pro-survival BCL-2 family proteins in OvCa. We therefore performed BH3 profiling on a panel of ten HGSOC cell lines, each with mutant p53^31^. All of the cell lines initiated mitochondrial apoptosis (as indicated by release of cytochrome c) in response to high doses (100 μM) of BIM and BID BH3 peptides (Figure 1A and S1A). These peptides are able to inhibit all pro-survival BCL-2 family proteins and also activate BAX and BAK directly^18,32^. Differences between these cell lines were evident at lower doses – the most primed of the cell lines were OVCAR8, OVCAR3 and OVKATE cells. Moderately primed cell lines included JHOM-1, KURAMOCHI, OAW28, OV90, OVCAR4 and TYK-nu cells. OVSAHO cells were least primed, as evidenced by their limited release of cytochrome c in response to the highest doses of pro-apoptotic BIM and BID BH3 peptides.

**Figure 1:**
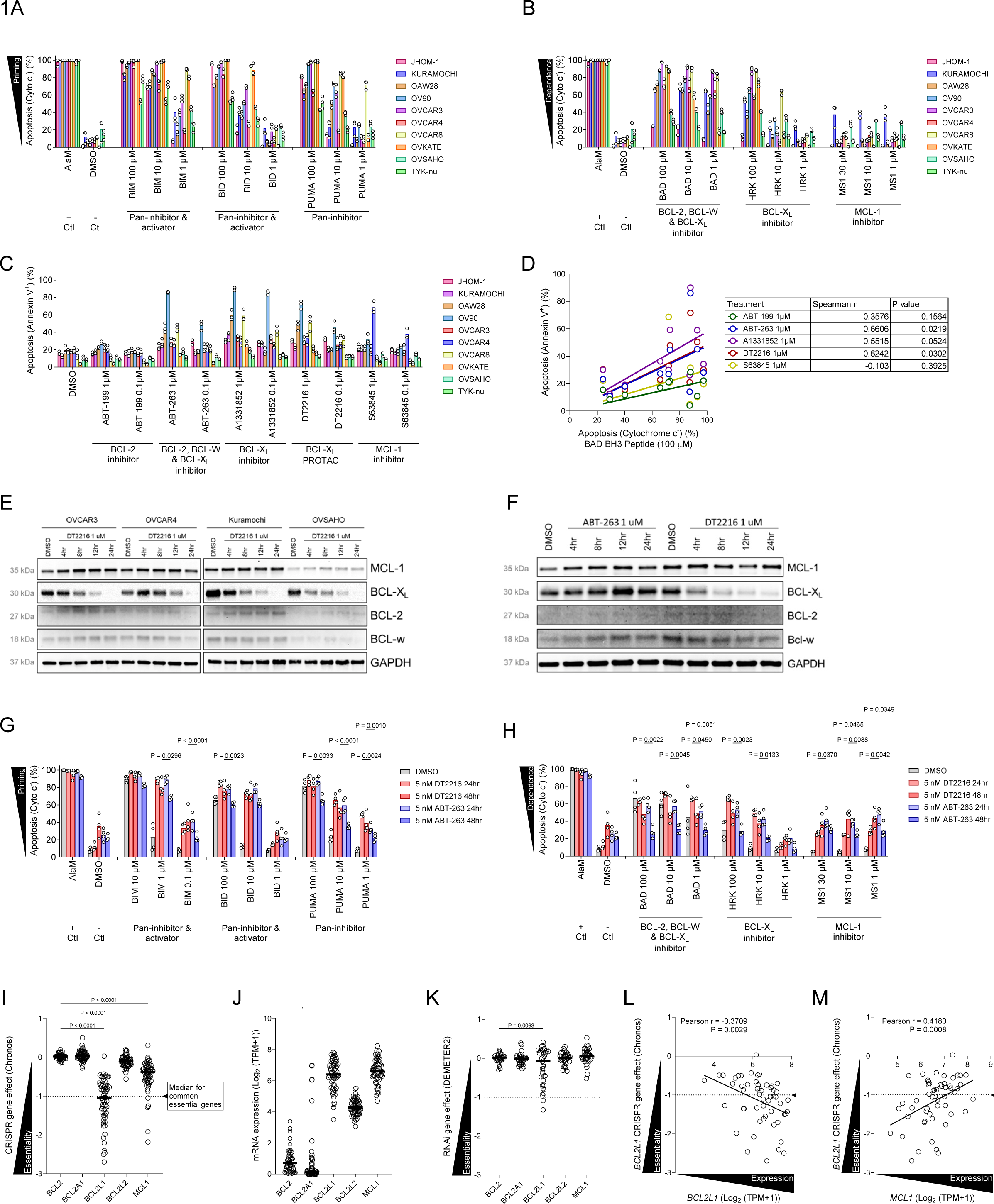
Ovarian cancer (OvCa) cell lines are primed for apoptosis and dependent on BCL-X_L_ for survival. (A) BH3 profiling of OvCa cell lines with BIM and BID activator peptides that inhibit all pro-survival proteins and also directly activate BAX and BAK and the PUMA BH3 sensitizer peptide that only inhibits all pro-survival proteins but does not activate BAX or BAK. Cytochrome c release, as indicated by the percentage of cells that are cytochrome c negative, signals initiation of apoptosis and indicates level of apoptotic priming. (B) BH3 profiling of OvCa cell lines with BAD, HRK and MS1 peptides that only inhibit indicated pro-survival proteins and do not activate BAX or BAK. Cytochrome c release indicates initiation of apoptosis and indicates level of dependence on pro-survival protein being inhibited. Mean ± SEM is shown for n = 3 biological replicates. (C) Annexin V/ PI staining and flow cytometry analysis of OvCa cell lines treated with agents targeting BCL-2 family proteins including ABT-199 (inhibits BCL-2 only), ABT-263 (inhibits BCL-2, BCL-X_L_ and BCL-W), DT2216 (degrades BCL-X_L_), A1331852 (inhibits BCL-X_L_), or S63845 (inhibits MCL-1) for 72 hours. Mean ± SEM is shown for n = 3 biological replicates. (D) Spearman’s rho correlation analysis comparing responses to BAD BH3 peptide at 100 μM in BH3 profiling versus apoptosis induced by indicated BH3 mimetic in annexin/PI chemosensitivity analysis. (E) Immunoblotting analysis of indicated proteins in OvCa cells treated with DT2216 for indicated duration. (F) Immunoblotting analysis of indicated proteins in OVCAR3 cells treated with ABT-263 or DT2216. (G-H) BH3 profiling of KURAMOCHI cells that were treated with DT2216 or ABT-263 for 24 or 48 hours to measure apoptotic priming (G) or dependencies (H). Mean is shown for n = 3 biological replicates. (I-M) Data from the Cancer Dependency Map (DepMap) showing (I) effect of CRISPR knockout of indicated gene on fitness of OvCa cell lines, (J) mRNA expression of pro-survival BCL-2 family genes, (K) effect of siRNA knockdown of indicated gene on fitness of OvCa cell lines, (L-M) comparison of the effects of BCL2L1 knockout and expression of BCL2L1 (L) or MCL1 (M) mRNA expression levels. Data in A-C are shown from individual experiments with bars representing means ± SEM from at least n=3 independent experiments. P values compare biological replicates in a one- or two-way ANOVA with Holm-Sidak’s adjustment for multiple comparisons.

These differences were consistent with the responses to the PUMA BH3 peptide, which can also inhibit all pro-survival proteins but cannot directly activate BAX and BAK^33^ – OVSAHO cells were again the most resistant to this pro-apoptotic BH3 peptide. These results indicated that most HGSOC cell lines maintain high sensitivity to pro-apoptotic signals and would be expected to trigger this form of cell death in response to stress or damage that produces pro-apoptotic signaling.

Given that OvCa cell lines were primed for apoptosis, we next sought to determine if these cell lines exhibit dependence on pro-survival proteins from the BCL-2 family. Using BH3 peptides that selectively inhibit each of these BCL-2 family pro-survival proteins, we detected consistent dependence on the pro-survival protein BCL-X_L_ in all OvCa cell lines except JHOM-1, OVSAHO and TYK-nu (Figure 1B and S1A). This was evident based on the sensitivity (cytochrome c release) of most OvCa cell lines to the BAD BH3 peptide that selectively inhibits BCL-2, BCL-X_L_ and BCL-W, as well as high sensitivity to the HRK BH3 peptide, which selectively inhibits BCL-X_L_. There was relatively low sensitivity to the MCL-1 inhibiting peptide MS1, demonstrating that this dependence is relatively weak and less consistently evident.

To further evaluate dependencies on pro-survival proteins, we tested OvCa cell line sensitivity to BH3 mimetics that selectively inhibit certain pro-survival proteins. Consistent with the BH3 profiling results, we detected that many HGSOC cell lines exhibited a loss of viability in response to BH3 mimetics that inhibit BCL-X_L_ including ABT-263, A1331852 and DT2216 but not the selective BCL-2 inhibitor ABT-199 or the selective MCL-1 inhibitor S63845 (except OVCAR4 cells) (Figure 1C). Furthermore, flow cytometric analysis of cells stained with the phosphatidylserine (PS) binding agent Annexin V (AxV) and DNA binding agent propidium iodide (PI) indicated that cells were undergoing apoptosis (Figure S1B). This is demonstrated by cells becoming positive for AxV prior to becoming positive for PI since the latter agent only enters cells with a disrupted plasma membrane, which occurs after externalization of PS in cells undergoing caspase-mediated apoptosis. When comparing the BH3 profiling with mimetic sensitivity analysis, Spearman’s rho correlational testing indicated that mitochondrial sensitivity to the BAD BH3 peptide correlated with cellular sensitivity to BH3 mimetics targeting BCL-X_L_ including ABT-263 and DT2216 (Figure 1D). This confirms that BH3 profiling can detect dependence on pro-survival proteins in ovarian cancer cells, as has been shown previously in other settings^17,34–37^.

We next sought to compare the apoptosis-promoting activity of ABT-263 and DT2216 since both of these agents have similar structures with only one major alteration: DT2216 includes a VHL ligand to facilitate BCL-X_L_ degradation^29^. We confirmed that the BCL-X_L_ degrading agent DT2216 rapidly and selectively reduces expression of BCL-X_L_ (Figure 1E) and triggers caspase-mediated, apoptotic cell death, which can be prevented by co-treatment with the pan-caspase inhibitor QVD-OPH (Figure S1C). Surprisingly, we also detected a very mild yet consistent increase in expression levels of pro-survival MCL-1 protein (Figure 1E), suggesting that some apoptotic adaptation may occur in response to BH3 mimetics. We directly compared this adaptation between ABT-263 and DT2216 and found that while mild upregulation of MCL-1 was evident in response to both agents, treatment with ABT-263 also caused upregulation of BCL-X_L_ to ostensibly buffer against induction of apoptosis (Figure 1F). This was completely absent in DT2216 treated cells due to the ongoing degradation facilitated by this agent, suggesting that the degree of apoptotic adaptation is limited by agents that degrade pro-survival proteins instead of inhibiting them. BH3 profiling of OvCa cells treated with ABT-263 or DT2216 demonstrated that although both agents increased apoptotic priming, ABT-263 treated cells exhibited a reduction in apoptotic priming and dependence on MCL-1 after 48 hours whereas DT2216 cells did not (Figure 1G). The enhanced apoptosis-promoting activity of DT2216 compared with ABT-263 was also evident in response to low-dose treatment with these agents for even longer periods (Figure S1D-E).

We next tested the extent to which these dependencies would be evident in an even broader set of OvCa cell lines. We compared the reduction of fitness across 58 OvCa cell lines in which each of the five major pro-survival genes are knocked out via CRISPR as part of the Broad Institute’s Cancer Dependency Map (DepMap) project^38^. None of the 58 cell lines were dependent on *BCL2, BCL2A1* (encoding A1) or *BCL2L2* (encoding BCL-W) for maintaining fitness (Figure 1I). Four of the cell lines (6.9%) exhibited a loss of fitness in response to knockout of *MCL1*, indicating that this gene is considered essential in a subset of OvCa cell lines. In contrast to the lack of widespread dependence on these pro-survival proteins, knockout of *BCL2L1* (encoding BCL-X_L_) resulted in a loss of fitness in 28 of 58 (48.3%) lines, indicating that this gene is essential for survival for about half of all OvCa cell lines (Figure 1I). The high dependence on this pro-survival protein in the CRISPR screen was consistent with the high expression of *BCL2L1* transcripts across OvCa cell lines (Figure 1J). Furthermore, a comparison of the effects on fitness in OvCa cell lines undergoing RNAi-mediated knockdown of pro-survival genes confirmed that only *BCL2L1* is an essential gene in the OvCa cell lines tested (Figure 1K). Consistent with the higher expression of *BCL2L1* in OvCa being an indicator of potential dependence, we found that mRNA expression of this gene is correlated with its essentiality (Figure 1L). Interestingly, high expression of *MCL1* transcripts was associated with reduced essentiality of *BCL2L1* (Figure 1M), suggesting that these proteins have some overlap in function and the overexpression of one can prevent dependency on the other.

### Ovarian primary tumors express BCL-X_L_, especially HGSOC

Since we found that high expression of *BCL2L1* transcripts is correlated with BCL-X_L_ dependence, we asked whether BCL-X_L_ protein was detectable in primary tumors. Using publicly available databases, we found that BCL-X_L_ is expressed in the majority of OvCa primary tumors available for analysis, including 6/6 HGSOC (Figure 2A-B). Half of HGSOC tumors expressed BCL-X_L_ in over 75% of cells with the remaining tumors expressing this protein in 25-75% of cells (Figure 2A, 2C). In all cases where BCL-X_L_ was expressed, including in mucinous, endometroid and serous OvCa, expression of the protein was found to be cytoplasmic or membranous (Figure 2A, 2D), consistent with the pro-survival effect that this protein has while interacting with other BCL-2 family proteins on the outer mitochondrial membrane.

**Figure 2:**
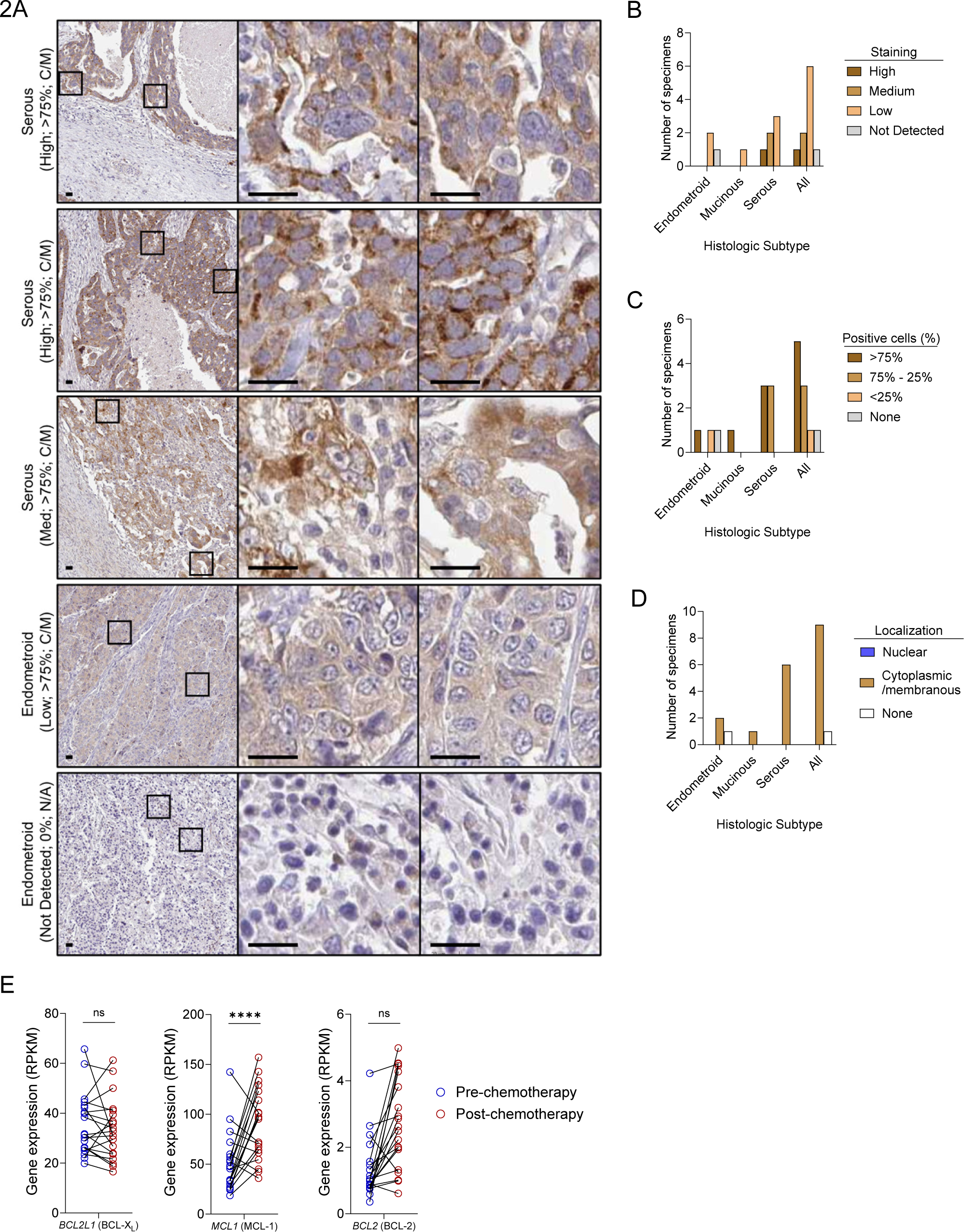
HGSOC tumors commonly express cytoplasmic BCL-X_L_. (A) Immunohistochemical analysis of BCL-XL expression across subtypes of OvCa tumors. Subtype, expression level (High, Medium, Low or Not Detected), % of cells staining positive and localization of BCL-X_L_ are noted for each specimen). (B-D) BCL-X_L_ staining analysis across subtypes of ovarian tumors. (E) Gene expression analysis from RNA-seq performed on HGSOC primary tumors before and after administration of neoadjuvant chemotherapy. P values compare biological replicates in a one-way ANOVA with Holm-Sidak’s adjustment for multiple comparisons.

We also sought to compare expression levels of BCL-2 family genes in primary HGSOC tumors and how they are altered by neoadjuvant therapy to understand how tumors evolve under therapeutic pressure. Analysis of an existing gene expression dataset from 20 patients with advanced stage (IIIC or IV) HGSOC prior to and after receiving neoadjuvant chemotherapy^39^ showed that pro-survival gene expression largely mirrored the OvCa cell line data (Figure 1J) with *BCL2L1*, *BCL2L2* and *MCL1* being consistently expressed at high levels (Figure S2A). Pro-apoptotic genes including *BAD* and *BCL2L11* (BIM) were consistently detected, as were the critical pore-forming proteins *BAX* and *BAK1* (Figure S2A). Interestingly, neoadjuvant therapy, which consisted of carboplatin and paclitaxel, did not significantly alter the expression of BCL-2 family genes in tumor cells except for MCL1, which was upregulated post neoadjuvant chemotherapy (Figure 2E and S2A). We also did not detect any significant associations between expression of specific BCL-2 family genes with outcomes in response to neoadjuvant chemotherapy (Figure S2B). Taken together, these data indicate that BCL-X_L_ is expected to be a consistently expressed and targetable apoptotic dependency in HGSOC. The increase in *MCL1* detected in post-therapy specimens may result in increased resistance to BCL-X_L_ inhibition, suggesting that treatment-naïve HGSOC tumors will be most susceptible to BCL-X_L_ inhibition.

### Dependence on BCL-X_L_ is enhanced by carboplatin and paclitaxel

Although our investigation demonstrated that BCL-X_L_ may be a targetable dependency in OvCa, a previous phase II trial testing the effects of ABT-263 in heavily pretreated women with recurrent epithelial OvCa showed that single-agent treatment is insufficient to produce remissions in the majority of patients^40^. We therefore sought to identify a combination therapy that would have more activity in the recurrent setting. We initially utilized the OVCAR3 cell line, which is a well-characterized model of chemoresistant and p53-mutant HGSOC^31^. Dynamic BH3 profiling of OVCAR3 cells treated with the standard of care agents carboplatin and paclitaxel showed that both of these agents increase apoptotic priming, indicating that treated cells are experiencing apoptotic stress (Figure 3A). Treatment with sensitizer BH3 peptides demonstrated that dependence on BCL-X_L_ is enhanced by paclitaxel treatment (Figure 3B).

**Figure 3:**
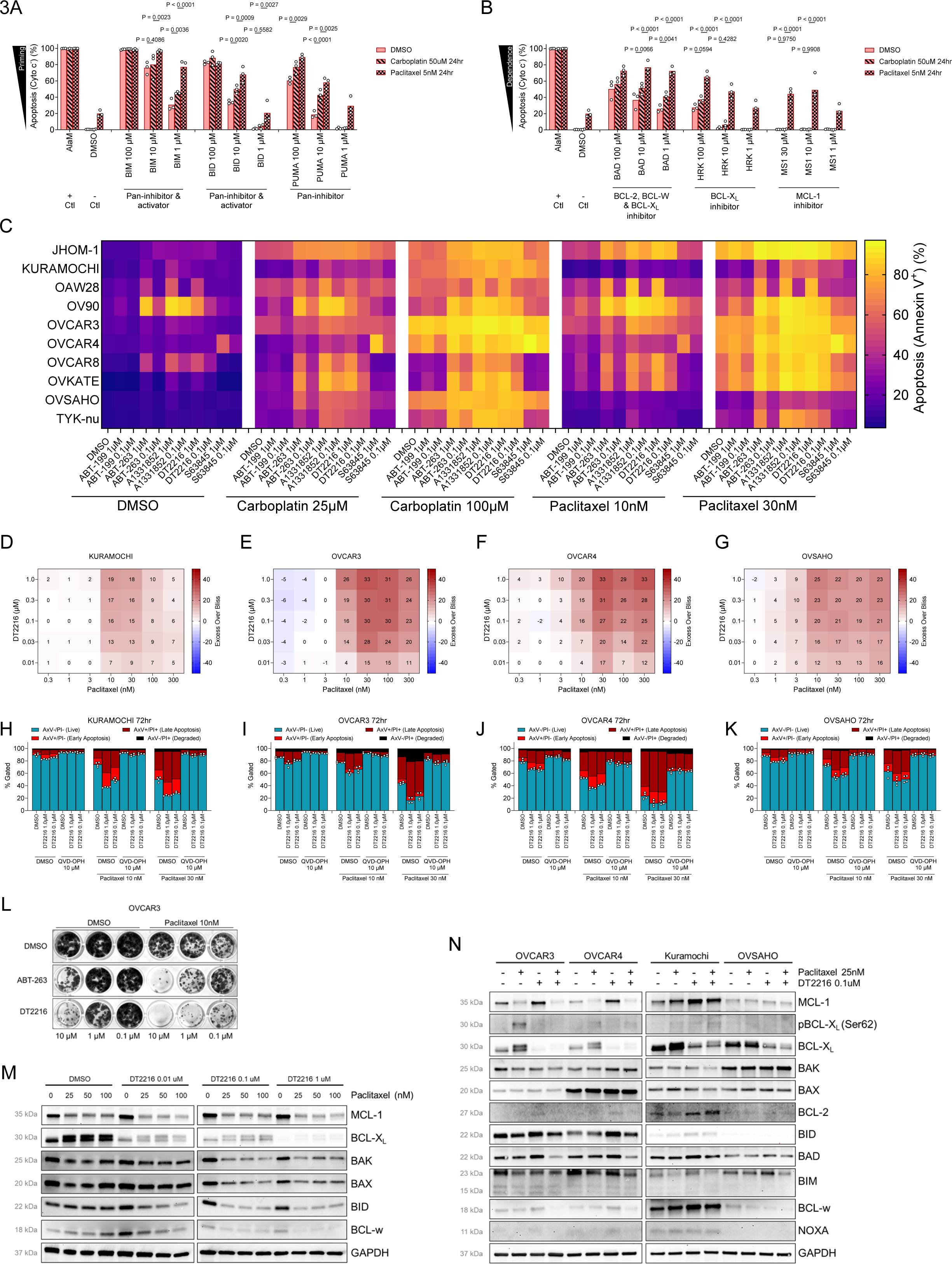
Paclitaxel and carboplatin increase apoptotic priming and BCL-X_L_ dependence in OvCa cell lines. (A-B) BH3 profiling of OvCa cell lines that were treated with carboplatin or paclitaxel for 24 hours to measure apoptotic priming (A) or dependencies (B). Mean ± SEM is shown for n = 4 biological replicates. (C) Annexin V staining and flow cytometry analysis of OvCa cell lines treated with carboplatin or paclitaxel in combination with agents targeting BCL-2 family proteins. Mean is shown for n = 3 biological replicates. (D-G) Bliss synergy analysis of OvCa cell lines treated with paclitaxel and DT2216. (H-K) Annexin V/ PI staining and flow cytometry analysis of OvCa cell lines treated with paclitaxel and DT2216 in the presence of pan-caspase inhibitor QVD-OPH for 72 hours. Mean ± SEM is shown for n = 4 biological replicates. (L) Colony formation assay performed on OVCAR3 cells treated with paclitaxel in combination with indicated agents targeting BCL-X_L_. Data are representative of n = 3 biological replicates. (M) Immunoblotting analysis of indicated proteins in OVCAR3 cells treated with indicated agents for 24 hours. (N) Immunoblotting analysis of indicated proteins in OvCa cell lines treated with indicated agents for 24 hours. P values compare biological replicates in a one- or two-way ANOVA with Holm-Sidak’s adjustment for multiple comparisons.

Based on the dynamic BH3 profiling results, we reasoned that inhibitors of BCL-X_L_ would enhance apoptosis induced by paclitaxel in OvCa cell lines more so than BH3 mimetics targeting other pro-survival proteins. As expected, we detected increased rates of apoptosis in HGSOC cell lines treated with each of the BCL-X_L_ inhibitors in combination with either standard of care agent, but particularly with paclitaxel (Figure 3C). Bliss independence analysis showed that the BCL-X_L_ PROTAC DT2216 synergized with paclitaxel across a range of concentrations, as indicated by “excess over Bliss” values in excess of 10^41^ (Figure 3D-G and S3A-D) and we confirmed that the cell death being induced was apoptosis using QVD-OPH (Figure 3H-K). Finally, colony formation assays showed that while single agent treatment with paclitaxel or DT2216 could reduce OVCAR3 colony growth, combined treatment with both agents completely abrogated the outgrowth of colonies (Figure 3L). Notably, an equivalent dose of DT2216 was more effective than ABT-263 at reducing colony growth as a single agent or in combination with paclitaxel at multiple concentrations, again suggesting that degradation of BCL-X_L_ may forestall the apoptotic adaptation that occurs in response to first-generation BCL-X_L_ inhibitors.

To gain insight into the mechanisms that are responsible for this synergistic apoptosis, we evaluated expression of BCL-2 family proteins in OVCAR3 cells treated with paclitaxel ± DT2216. Consistent with our previous results, we found that protein expression levels of BCL-X_L_ were reduced in a dose-dependent manner by DT2216 (Figure 3M). We also detected a consistent increase in MCL-1 expression in OVCAR3 cells treated with DT2216 at three different concentrations (Figure 3M). This was also evident in the other cell lines treated with DT2216 (Figure 3N). In contrast, paclitaxel treatment led to a decrease in MCL-1 protein expression, which would be expected to lead to the increased priming that we detected. Interestingly, we also detected a second, higher-weight protein band in the immunoblot for BCL-X_L_ in cells treated with paclitaxel (Figure 3M-N), which we reasoned may be indicative of a phosphorylation event. BCL-X_L_ phosphorylation at Ser62 has been previously found to inactivate the pro-survival activity of this protein and is likely also contributing to the increased priming and BCL-X_L_ dependence we detected in paclitaxel-treated cells (Figure 3A-B). Using a phospho-specific antibody, we detected phosphorylation of BCL-X_L_ at Ser62 in paclitaxel-treated OVCAR3 cells (Figure 3N), confirming that this inactivating event is occurring in response to microtubule targeting with this agent.

### Ovarian primary tumors are dependent on BCL-X_L_ for survival

Cancer cell lines can accumulate alterations during in vitro culture that limit their effectiveness as disease models^31,42,43^. To address this, we performed BH3 profiling, which does not require any ex vivo culture, and chemosensitivity analysis on seven freshly-collected HGSOC primary tumors from different disease phases of treatment immediately after tumor resection (Table 1). HGSOC cells from each of the seven specimens were highly responsive to pro-apoptotic BH3 peptides even at low concentrations (BIM BH3 peptide at 1 μM, for example), indicating that HGSOC primary tumors can readily initiate apoptosis in response to pro-apoptotic signaling, even at moderate or low levels (Figure 4A). We noted that even tumor specimens from patients with recurrent disease continued to be primed for apoptosis – this suggests that targeting the apoptosis pathway would be effective even at relapse after initial therapy. Next, we evaluated responses to the sensitizer peptides that can indicate dependence on individual BCL-2 family pro-survival proteins. We detected cytochrome c release in response to the BAD BH3 peptide (100 μM) that inhibits BCL-2, BCL-W and BCL-X_L_ in 6 of 7 primary OvCa specimens based on comparisons to DMSO treatment (Figure 4B). We detected BCL-X_L_ dependence using the HRK BH3 peptide (100 μM) in all seven HGSOC primary tumors, including treatment naïve, neoadjuvant and recurrent patients. We also detected dependence on MCL-1 in five of seven HGSOC specimens (Figure 4B) based on responses to the MS1 peptide (30 μM). Consistent with the HGSOC cell line data, using dynamic BH3 profiling on HGSOC specimen DF4379 we found that paclitaxel treatment increased primary OvCa cell dependence on BCL-X_L_, as indicated by increased cytochrome c release in response to the BAD and HRK BH3 peptides (Figure 4C-D).

**Figure 4:**
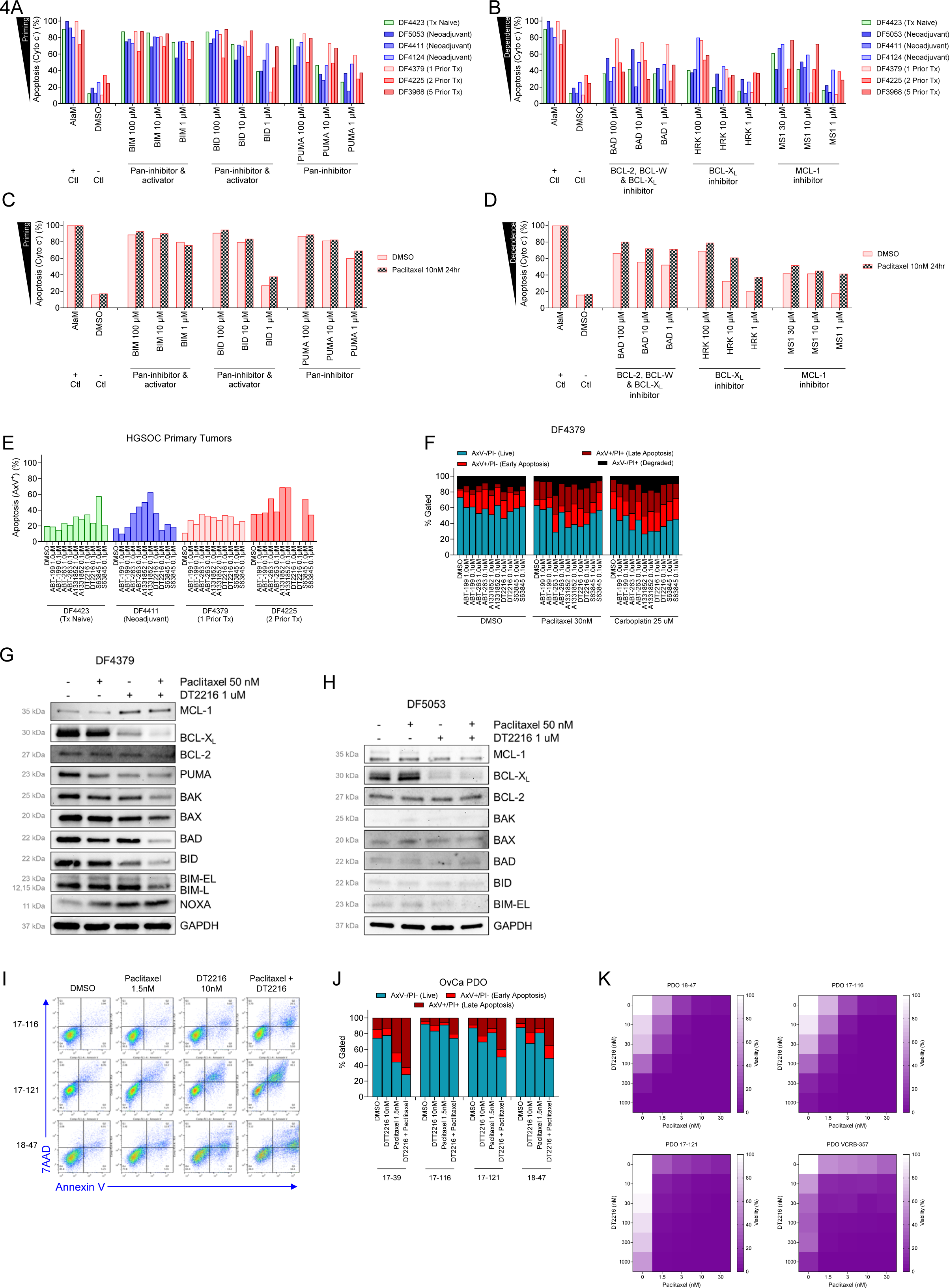
OvCa primary tumors and patient-derived organoids are primed for apoptosis and dependent on BCL-X_L_. (A-B) BH3 profiling of OvCa primary tumors to measure apoptotic priming (A) or dependencies (B). (C-D) BH3 profiling of OvCa primary tumor DF4379 treated with carboplatin or paclitaxel for 24 hours to measure apoptotic priming (C) or dependencies (D). Mean is shown for n = 1-2 technical replicates. (E) Annexin/ PI staining and flow cytometry analysis of OvCa primary tumors treated with agents targeting BCL-2 family proteins. Mean is shown for n = 1-2 technical replicates. (F) Annexin/PI staining and flow cytometry analysis of OvCa primary tumor DF4379 treated with paclitaxel and agents targeting BCL-2 family proteins for 72 hours. Mean is shown for n = 1-2 technical replicates. (G-H) Immunoblotting analysis of DF4379 (G) and DF5053 (H) treated with indicated agent for 24 hours. (I-J) Annexin/7AAD staining and flow cytometry analysis of OvCa patient derived organoids treated with paclitaxel and DT2216 for 5 days. Mean is shown for n = 1 biological replicates. (K) ATP measurements in OvCa patient derived organoids treated with paclitaxel and DT2216 for 5 days.

Ex vivo chemosensitivity analysis of the freshly-collected HGSOC primary tumors was consistent with BH3 profiling data and indicated that BCL-X_L_ inhibition with single-agent treatment with A1331852, ABT-263 or DT2216 induced apoptosis (Figure 4E), as well as inhibition of MCL-1 with S63845 in certain tumors. We also confirmed that combination treatment with paclitaxel and BCL-X_L_ inhibitors more potently induced apoptosis in DF4379 (Figure 4F), consistent with the dynamic BH3 profiling results (Figure 4D).

We next evaluated BCL-2 family protein changes in HGSOC primary tumors in response to paclitaxel and DT2216 treatment ex vivo. We confirmed that DT2216 treatment led to the rapid and preferential degradation of BCL-X_L_ in primary HSGOC tumor cells (Figure 4G), which was further enhanced in the presence of paclitaxel. Of note, we again detected a second band in the BCL-X_L_ immunoblots, even in the absence of ex vivo treatment with paclitaxel. Similarly, we also noted an increase in MCL-1 expression in response to DT2216 treatment, suggesting the potential for rapid adaptation to this treatment. Co-treatment with paclitaxel again mitigated this effect, showing that combination treatment may prevent any rapid adaptations that could confer resistance to BH3 mimetic treatment. Similar effects, including the rapid degradation of BCL-X_L_, by DT2216 and paclitaxel-mediated phosphorylation of this protein were observed in the DF5053 specimen (Figure 4H).

### Ovarian patient-derived organoids are dependent on BCL-X_L_ for survival after therapy-induced apoptotic stress

Patient-derived organoids (PDOs) provide a platform for assessment of tumor cell sensitivity to therapy in tumor cells grown as spheroids, which more accurately reflects the morphology of patient tumors and can be more predictive of in vivo therapy efficacy^44^. We tested the sensitivity of five HGSOC PDO models that reflect different phases of disease, from treatment-naïve to recurrent (Table 2), to combination therapy including paclitaxel and DT2216 and assessed death by quantifying apoptosis via AxV/7AAD staining. The treatment-naïve PDO, 18-47, was mildly sensitive to paclitaxel treatment and exhibited more sensitivity to low-dose treatment with DT2216 (Figure 4I-J). Impressively, the combination treatment was most effective, and induced more apoptosis than would be expected based on additivity alone (Figure 4J). The neoadjuvant treated PDO, 17-116, exhibited very limited sensitivity to either of the agents alone but again the combination treatment resulted in enhanced apoptosis. The two heavily pretreated, recurrent samples exhibited some divergence in sensitivity: 17-39 exhibited no sensitivity to DT2216 alone but, when DT2216 was combined with paclitaxel, there was enhancement of apoptosis. Finally, the 17-121 specimen exhibited very limited sensitivity to paclitaxel while being sensitive to single-agent DT2216 and even more sensitive to combination treatment. These tests of apoptotic cell death were consistent with results obtained via ATP measurement (Figure 4K). Overall, these data demonstrated that HGSOC tumor cells, when grown in tumor-like organoids, continued to exhibit strong dependence on BCL-X_L_, especially in combination with paclitaxel treatment.

### Ovarian PDX models are dependent on BCL-X_L_, sensitive to DT2216

Short-term ex vivo culture of OvCa PDX models allows for rapid evaluation of therapeutic efficacy across a diverse range of HGSOC models that exhibit immunohistological and molecular fidelity to the original patient tumor^45^. We collected ascites from ten luciferized ovarian cancer PDX models (Table 3) generated from patients with varying clinical and genetic characteristics including prior treatment history (ranging from treatment-naïve to 7 prior rounds of therapy), BRCA status (WT as well as BRCA1 mutant), and HGSOC-associated mutations (e.g., *TP53, PTEN, CDKN2A*) that were growing in immunocompromised mice. These OvCa cells were grown in vitro for two days, treated with DT2216 and paclitaxel for four days and then cell viability was evaluated by adding luciferin and measuring luminescence. We noted that treatment naïve (e.g., DF09) as well as heavily pretreated PDX models (e.g., DF14 – 5 prior chemo cycles; DF181 – 7 prior cycles) were highly sensitive to DT2216 treatment alone (Figure 5A). This suggests that HGSOC cells are dependent on BCL-X_L_ regardless of prior treatment status, which is in agreement with our RNA-seq data (Figure 2E and S2A). The results in DF14 and DF181 were particularly noteworthy given their resistance to paclitaxel treatment, presumably due to their extensive prior treatment with this agent. In total, 70% of PDX models exhibited sensitivity to single agent DT2216, as defined by 33% or greater loss of viability in response to DT2216 treatment at 0.71 μM. We also noted that 6/10 PDX models were more sensitive to DT2216 than paclitaxel at equivalent doses. Finally, we detected synergy between DT2216 and paclitaxel treatment in 7/10 PDX models.

**Figure 5:**
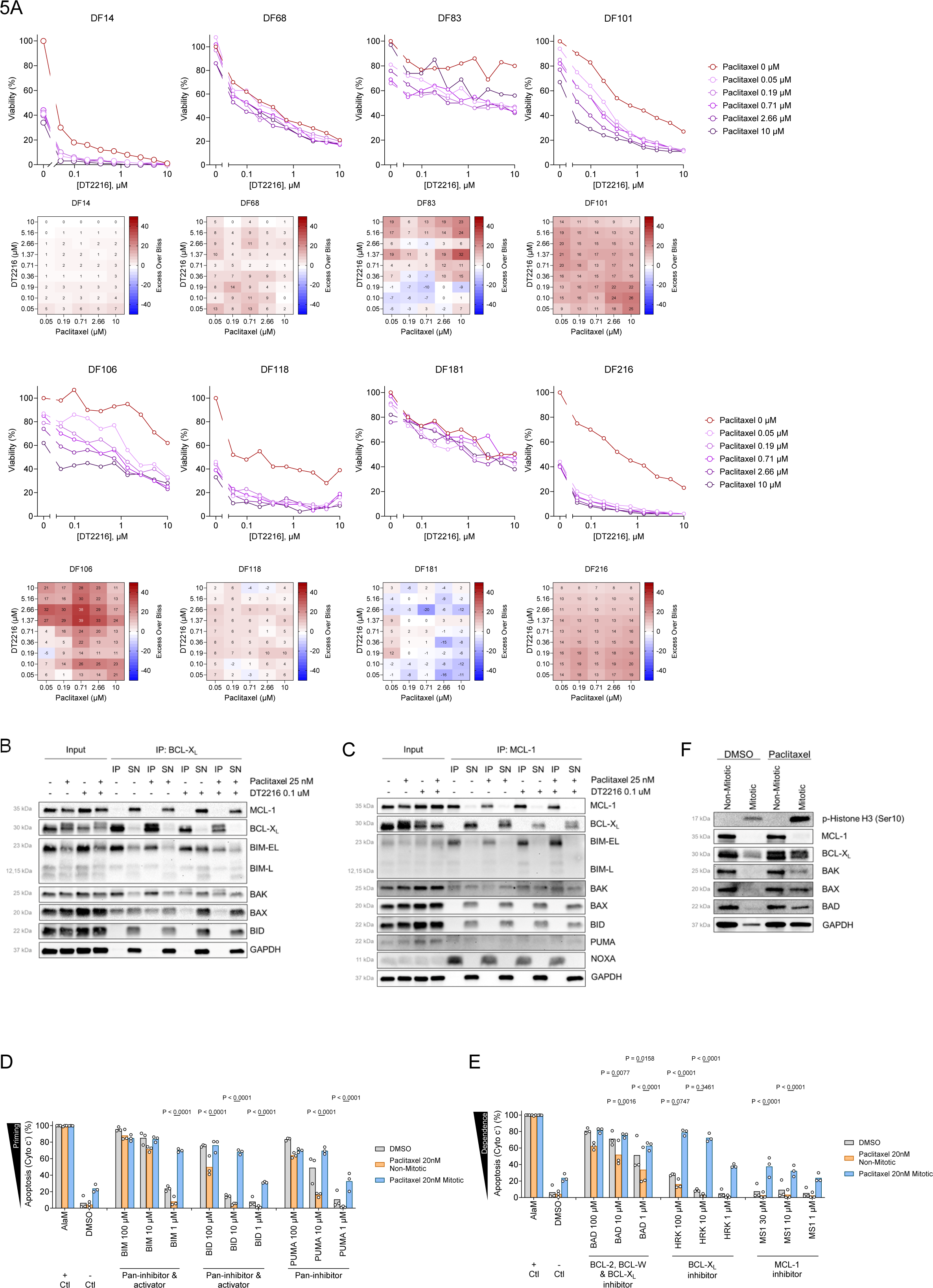
HGSOC PDX models are broadly sensitive to combination treatment with paclitaxel and DT2216 due to release of BIM, BAX and BAK from BCL-X_L_. (A) OvCa PDX models collected during in vivo growth were cultured ex vivo for 48 hours, then treated with DT2216 ± paclitaxel and viability was measured after 96 hours by adding luciferin and measuring luminescence. Mean is shown for n = 4 technical replicates from one biological replicate that is representative of n = 2 biological replicates. (B-C) Immunoprecipitation of BCL-X_L_ (B) or MCL-1 (C) in OVCAR3 cells treated with DT2216 ± paclitaxel for 24 hours. Input lanes show all proteins collected in various treatments. IP indicates immunoprecipitation (bound proteins) while SN indicates supernatant (unbound proteins). Data are representative of n = 2 biological replicates. (D-E) BH3 profiling of non-mitotic and mitotic OVCAR3 cells that were treated with paclitaxel for 24 hours to measure apoptotic priming (D) or dependencies (E). Mean is shown for n = 3 biological replicates. (F) Immunoblotting of non-mitotic and mitotic OVCAR3 cells that were treated with paclitaxel at 20 nM for 24 hours.

### BCL-X_L_ restrains BAX, BAK and BIM in HGSOC cells

The consistent synergy between paclitaxel and DT2216 prompted us to investigate the binding interactions that lead to induction of apoptosis in response to this combination treatment. By immunoprecipitating BCL-X_L_ and blotting for BCL-2 family proteins in OVCAR3 cells, we detected active binding pro-apoptotic proteins including BIM, BAK and BAX, indicating active sequestration of multiple proteins that are critical for initiation of apoptosis (Figure 5B). In cells treated with paclitaxel, BCL-X_L_ continued to bind these pro-apoptotic proteins while levels of MCL-1 were reduced, contributing to increased BCL-X_L_ dependence. Low-dose DT2216 treatment led to BCL-X_L_ degradation, which freed the majority of bound BIM, BAK and BAX. Combination treatment further exacerbated these changes, with only minimal BIM remaining bound to BCL-X_L_. Immunoprecipitation of MCL-1 under these same conditions clarified the mechanism of apoptosis induction. In untreated cells, MCL-1 actively bound BIM and BAK but not BAX (Figure 5C). Treatment with paclitaxel reduced the expression of MCL-1 while the remaining protein continued to bind BIM and BAK, again contributing to increased priming and dependence on both BCL-X_L_ and MCL-1. Treatment with DT2216 forced MCL-1 to bind and sequester additional BIM that was freed from the degraded BCL-X_L_. Combination treatment causes a reduction in MCL-1 levels along with reduced ability for BCL-X_L_ to sequester the freed pro-apoptotic signals, thus triggering apoptosis.

### Prolonged mitotic arrest is associated with increased BCL-X_L_ dependence

Since paclitaxel is known to disrupt microtubule dynamics and cause mitotic arrest in cancer cells, we next sought to compare the levels of apoptotic priming and dependencies in cycling versus arrested cells. We first confirmed that paclitaxel treatment causes mitotic arrest of OvCa cells (Figure S5C) and then subjected the paclitaxel-treated cells to BH3 profiling. Notably, we detected a major difference in priming between cell populations: in the presence of paclitaxel, non-mitotic cells exhibited lower levels of priming than untreated cells while those cells that were undergoing mitosis were much more primed (Figure 5D). We also detected reduced BCL-X_L_ dependence in the non-mitotic cells while those cells that were arrested in mitosis were much more dependent on BCL-X_L_ and, to a lesser extent, MCL-1 (Figure 5E).

Immunoblotting analysis uncovered the likely molecular mechanism driving these changes: expression of MCL-1 is nearly absent in cells that are arrested in mitosis as a result of paclitaxel treatment while BCL-X_L_ levels are largely maintained (Figure 5F). Taken together, these data indicate that prolonged mitotic arrest is responsible for driving reduced MCL-1 expression and convergence of pro-survival dependencies onto BCL-X_L_, setting the stage for potent induction of apoptosis in response to BCL-X_L_ degradation with DT2216.

### Ovarian xenograft tumors are eradicated with combination treatment targeting BCL-X_L_ and microtubules

We next set out to test the efficacy and safety of combination therapy targeting microtubules with paclitaxel with BCL-X_L_ inhibition/degradation in vivo. The on-target and dose-limiting thrombocytopenia induced by small molecule inhibitors of BCL-X_L_ has prohibited the clinical utilization of these agents^46^. To reduce the likelihood of this issue precluding the clinical evaluation of the synergistic anti-OvCa activity of BCL-X_L_ inhibition in combination with paclitaxel, we compared the sensitivity of mouse and human platelets to BH3 mimetics targeting BCL-2 family proteins in vitro. We found that mouse platelets were largely insensitive to inhibitors of BCL-2 or MCL-1 but were highly sensitive to inhibitors of BCL-X_L_ except the BCL-X_L_ PROTAC DT2216, which did not affect mouse platelet viability (Figure 6A).

**Figure 6:**
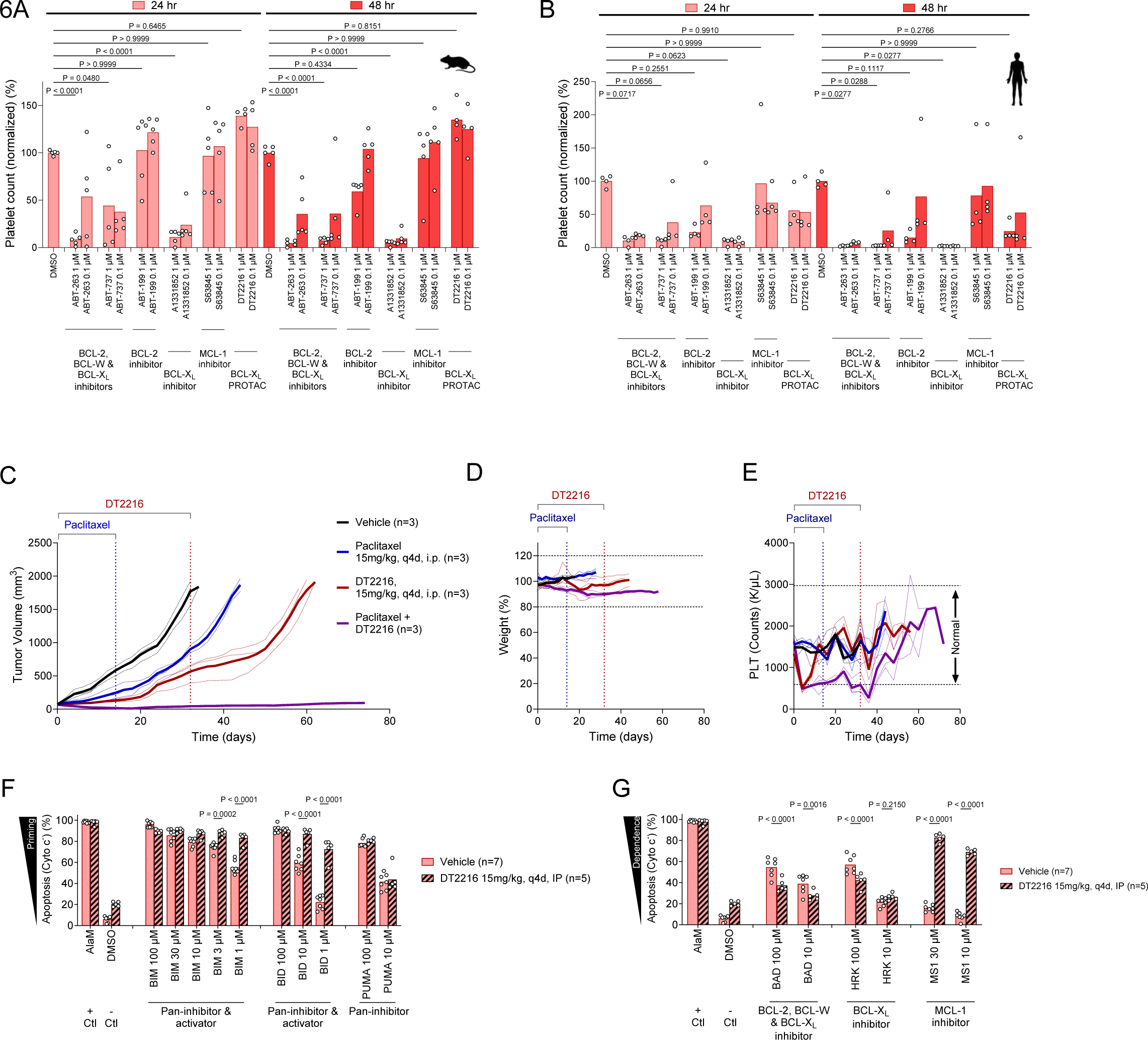
HGSOC xenograft tumors are sensitive to combination treatment with paclitaxel and DT2216. (A-B) Annexin/PI staining and flow cytometry analysis of platelets collected from mice (A) or humans (B) and treated in vitro with indicated BH3 mimetic or PROTAC. Mean is shown for n = 3 biological replicates. (C) OVCAR3 xenograft tumor growth in mice treated with indicated agents for indicated periods. (D-E) Weight (D) and platelet (E) measurements of mice treated with indicated agents. (F-G) BH3 profiling of OVCAR3 xenograft tumors at time of euthanasia treated with indicated agents to measure apoptotic priming (F) or dependencies (G). Mean is shown for n = 5 biological replicates. P values compare biological replicates in a one- or two-way ANOVA with Holm-Sidak’s adjustment for multiple comparisons.

These results were similar to analyses performed on human platelets (Figure 6B). These data indicated that DT2216 is less toxic to platelets than the first-generation small molecule BCL-X_L_ inhibitors.

To evaluate the efficacy of the BCL-X_L_ targeting therapy we identified, we established xenografts of the OVCAR3 cell line, which has been a well-validated model of aggressive HGSOC^47^, in immunocompromised NSG (NOD.Cg-*Prkdc^scid^ Il2rg^tm1Wjl^*/SzJ) mice. Once tumors reached a size of about 70-100 mm^3^, we administered paclitaxel at 15 mg/kg every four days via intraperitoneal injection for the first 14 days, with a total dose of 75 mg/kg per animal, DT2216 at 15 mg/kg every four days, or a combination of these treatments. These tumors were largely insensitive to paclitaxel treatment, which only slowed tumor growth (Figure 6C) – this is consistent with OVCAR3 resistance to this agent in vitro (Figure 3D). DT2216 was more effective as a single agent therapy and slowed tumor growth further but did not induce tumor remissions (Figure 6C). Impressively, the combination treatment caused near-complete tumor regression with no detected tumor outgrowth even after cessation of therapy (Figure 6C).

To monitor for signs of toxicity, we monitored animal weight and performed complete blood counts every four days. We detected, on average, less than 10% body weight loss in animals treated with any of the agents including the combination (Figure 6D). Due to the on-target thrombocytopenia that is caused by inhibition of BCL-X_L_ in patients, we closely monitored platelet counts and found that DT2216 caused only a detectable reduction in platelet levels after the first dose (Figure 6E). Platelet levels recovered to a normal range for all subsequent time points despite continuous DT2216 dosing. In this experiment, while the paclitaxel did not induce any reduction in platelet counts, the combination treatment of DT2216 with paclitaxel did delay platelet count recovery, which was achieved when DT2216 treatment concluded on day 32.

Further, in animals treated with DT2216, we performed BH3 profiling of tumors at time of euthanasia to understand how this treatment affects apoptosis sensitivity in tumor cells. Despite treatment having ceased over 30 days prior to tumor collection, we detected higher levels of apoptotic priming in DT2216 treated tumors compared with vehicle treated controls (Figure 6F). Further, we detected reduced dependence on BCL-X_L_ in these tumors and, surprisingly, enhanced dependence on MCL-1 (Figure 6G), suggesting that DT2216 treated tumors had adapted to BCL-X_L_ inhibition by upregulating MCL-1, which can be attenuated by paclitaxel.

### DT2216 eradicates HGSOC patient-derived xenograft (PDX) tumors in combination with paclitaxel

Finally, we sought to test the activity of paclitaxel and DT2216 in OvCa PDX tumors, which more faithfully recapitulate patient tumors than cell line-based xenografts. We established five OvCa PDX models originating from patients at various phases of disease (Table 3) in vivo to assess their levels of priming and BCL-X_L_ dependence. BH3 profiling of PDX tumor cells indicated that 4/5 of the PDX models were primed for apoptosis, again indicated by the high degree of cytochrome c release in response to even moderate (10 μM) or low (1 μM) doses of pro-apoptotic BIM and BID BH3 peptides (Figure 7A). Interestingly, the PDX model that was derived from the patient with the highest number of prior chemotherapy cycles (DF68 – 5 prior cycles of therapy) was also the least primed. We also detected significant release of cytochrome c in response to the BAD and HRK BH3 peptides in 3/5 PDX models, indicating detectable levels of dependence on BCL-X_L_ (Figure 7B).

**Figure 7:**
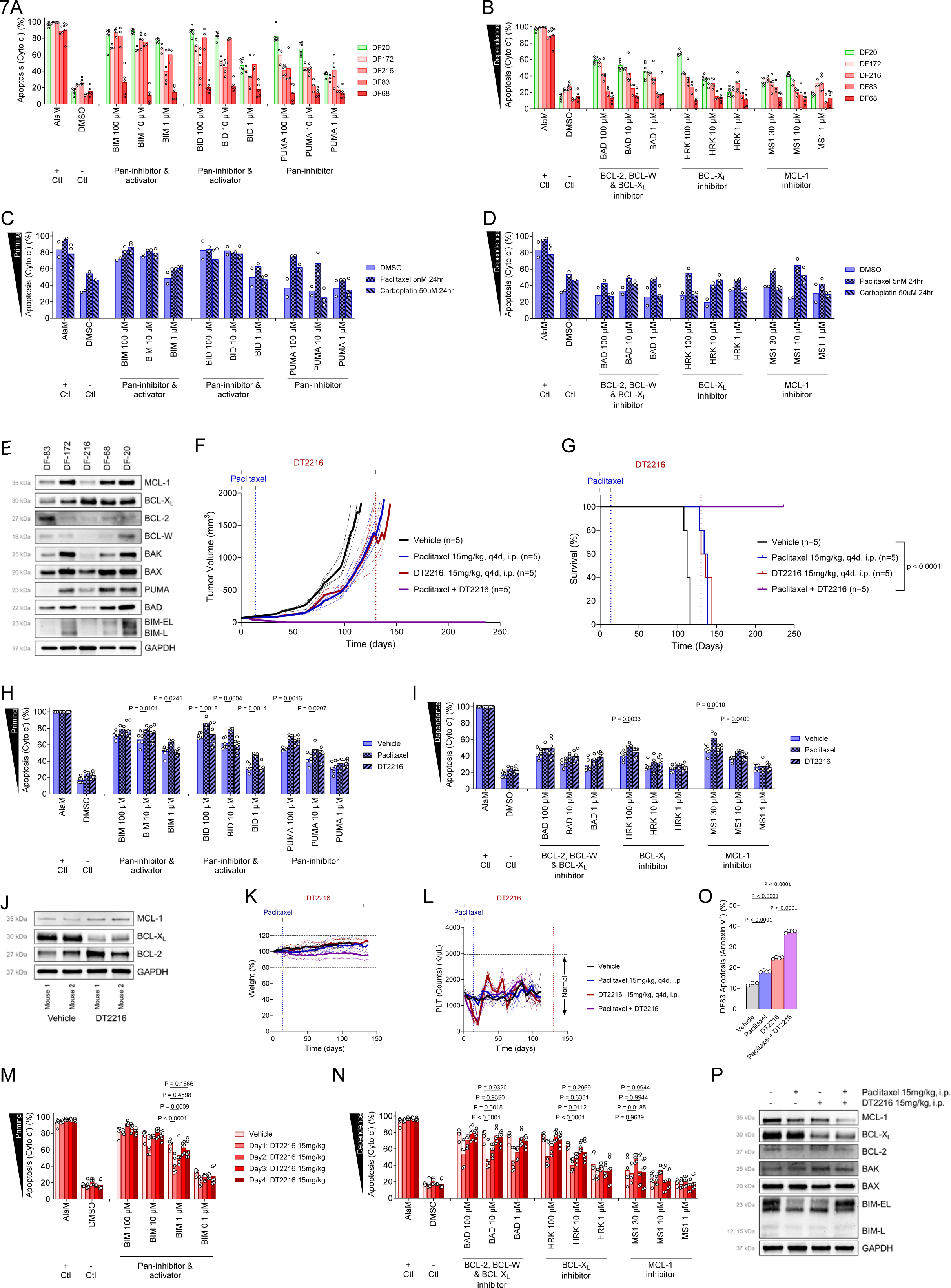
HGSOC patient-derived xenograft (PDX) tumors are sensitive to combination treatment with paclitaxel and DT2216. (A-B) BH3 profiling of OvCa PDX models to measure apoptotic priming (A) or dependencies (B). Mean is shown for n = 3-6 biological replicates. (C-D) BH3 profiling of DF83 PDX model during in vitro treatment with paclitaxel or carboplatin to measure apoptotic priming (C) or dependencies (D). Mean is shown for n = 3 biological replicates. (E) Immunoblotting analysis of indicated proteins in OvCa PDX models. (F) DF83 PDX tumor growth and survival of mice treated with indicated agents for indicated periods. (G-H) BH3 profiling of DF83 PDX tumors at time of euthanasia treated with indicated agents to measure apoptotic priming (G) or dependencies (H). Mean is shown for n = 3-5 biological replicates. (I) Immunoblotting of indicated proteins in DF83 PDX tumors at time of euthanasia treated with indicated agents. (J-K) Weight (J) and platelet counts (K) of mice treated with indicated agents. (L-M) BH3 profiling of platelets isolated from mice treated with indicated agents to measure apoptotic priming (L) or dependencies (M). Mean is shown for n = 7 biological replicates. (N) Annexin staining and quantification in EPCAM^+^ cells from DF83 xenograft tumor treated for 48 hours with indicated agents. Mean is shown for n = 4 biological replicates. (O) Immunoblotting analysis of indicated proteins in DF83 PDX tumors treated with indicated agents for 48 hours. Data in (A-D), (G-H) and (L-M) are shown from independent experiments with bars representing means from at least three independent experiments. P values were calculated using one- or two-way ANOVA with Holm-Sidak’s adjustment for multiple comparisons.

Although the DF20 PDX model was the most highly primed and most dependent on BCL-X_L_, this model was derived from a patient that was treatment naive. We intended to test the combination therapy in a recurrent model to determine whether this combination would be effective in early clinical trials that are most likely to include patients with recurrent disease. We therefore performed dynamic BH3 profiling on the recurrent DF83 model and detected significant increases in apoptotic priming (Figure 7C) and BCL-X_L_ dependence (Figure 7D) when cells were treated with paclitaxel. We found that the DF83 tumor cells expressed BCL-X_L_, albeit at lower levels than other models that had been treated with fewer cycles of chemotherapy (Figure 7E).

We established growth of DF83 PDX tumors and initiated a similar treatment regimen as used for OVCAR3 xenografts. Consistent with the prior treatment of these cells with chemotherapy, paclitaxel treatment only slowed tumor growth in vivo (Figure 7F). DT2216 also had a modest impact on tumor growth, consistent with potential adaptation to single-agent treatment. Strikingly, the combination of paclitaxel and DT2216 induced rapid tumor regressions and eradicated the tumors completely (Figure 7F). Furthermore, tumors showed no evidence of regrowth even more than 100 days after the cessation of DT2216 treatment (Figure 7G).

BH3 profiling of tumors treated with paclitaxel in vivo demonstrated a detectable increase in apoptotic priming (Figure 7H) and BCL-X_L_ dependence (Figure 7I), which supported the increased sensitivity of the cells to DT2216. Immunoblot analysis of DT2216 treated tumors uncovered an increase in expression of MCL-1 as well as BCL-2 (Figure 7J), suggesting that adaptation was occurring in response to DT2216, which did potently reduce levels of BCL-X_L_.

Toxicity assessment showed that these therapies were well tolerated, with no loss of body weight and only transient reduction in platelet counts (Figure 7K-L). Pathological examination of tissue samples from mice treated with vehicle, DT2216, paclitaxel, or a combination of both drugs revealed no detectable signs of drug toxicity in the spleen, lung, liver, kidney, or heart (Figure S7A). To gain further insight into the resiliency of platelets to DT2216 treatment, we BH3 profiled platelets in mice each day between treatments with DT2216. Interestingly, we detected reduced priming in platelets 1 and 2 days after dosing DT2216 as compared with vehicle-treated mice (Figure 7M) and also reduced dependence on BCL-X_L_ (Figure 7N). Interestingly, platelet dependence on MCL-1 was found to be elevated during this same period (Figure 7N). Over the course of the next three days, these measures returned to baseline levels comparable to vehicle treated mice. Since previous studies found that BCL-X_L_ inhibitor treatment leads to platelet loss and a compensatory increase in platelet production from megakaryocytes^48^, these results are consistent with platelets recently released into circulation being less primed for apoptosis and less dependent on BCL-X_L_ but more dependent on MCL-1.

In a separate cohort of DF83 PDX-bearing mice, we performed tumor cell analysis after short term (48 hour) treatment. We detected an enhancement of apoptosis induction in EPCAM+ DF83 tumor cells in combination treated mice as compared to single agents (Figure 7O). We also detected on-target degradation of BCL-X_L_ upon DT2216 treatment and decreased MCL-1 expression with paclitaxel treatment (Figure 7P). Taken together, these data indicate that combination treatment is most effective at inducing the changes in BCL-2 family protein expression and interactions that facilitate meaningful HGSOC tumor regressions.

## Discussion

Despite the increasing and well-founded interest in developing targeted therapies for a wide range of malignancies, cytotoxic chemotherapies have been proven to be highly effective and have undeniably led to the eradication of cancers in millions of patients, including women with HGSOC^2^. These chemotherapeutics, which include the HGSOC first-line therapies carboplatin and paclitaxel, target ubiquitous cell features, including DNA and microtubules, and typically kill cancer cells via the apoptosis pathway^10,17,49^. It is therefore important to identify apoptosis-promoting agents that can be used in combination with cytotoxic chemotherapies to enhance the chemosensitivity of cancer cells while maintaining a tolerable safety profile. Indeed, herein we have reported that BCL-X_L_ PROTAC degrader DT2216 can be safely combined with paclitaxel to induce extensive apoptosis in vitro and in vivo. In fact, this combination consistently eradicates HGSOC cell line and patient-derived xenograft tumors in vivo – a remarkable result given the inherent aggressiveness and chemoresistance of these models. Importantly, DT2216 induces only transient thrombocytopenia and no other detectable signs of toxicity, which suggests that this agent will be better tolerated by patients than first-generation inhibitors of BCL-X_L_. The lack of additional toxicity is consistent with our previous work showing that parenchymal cells within most adult healthy tissues are resistant to apoptosis due to low expression of pro-apoptotic BAX and BAK^50^, rendering them resistant to BCL-X_L_ inhibition^51^. We believe this is the first clinically viable combination strategy to effectively treat drug resistant OvCa by simultaneously targeting two of the most important pro-survival proteins selectively in tumor cells but not in normal tissues, i.e. direct degradation of BCL-X_L_ with DT2216 and indirect inhibition of MCL-1 with paclitaxel-induced mitotic arrest. This unique combination allows us to overcome the on-target toxicities caused by simultaneously targeting BCL-X_L_ and MCL-1 using direct small molecule inhibitors with or without conventional chemotherapy as shown in many previous studies. These toxicities include BCL-X_L_ inhibitor-induced dose-limiting thrombocytopenia^52,53^ and MCL-1 induced cardiotoxicity^54^. Further, combined direct inhibition of BCL-X_L_ and MCL-1 leads to severe and widespread tissue toxicity and lethality^51^. These on-target and dose-limiting toxicities have stalled the clinical development of these otherwise promising inhibitors. Our combination strategy avoids these toxicities and also produced a remarkable synergistic induction of apoptosis in OvCa cells, resulting in complete eradications of tumors and cures in both HGSOC cell line and patient-derived xenograft models without any detectable toxicities. That is remarkable given the inherent therapy resistance of the heavily pretreated PDX models that we have used for our studies^55^. Our finding has prompted the initiation of a phase Ib/II study using this combination to treat relapsed/refractory ovarian cancer patients.

Our BH3 profiling analysis of platelets after treatment with DT2216 provides novel insights into the regulation of apoptosis in these cells in a preclinical therapeutic setting. Specifically, we found that dosing mice with DT2216 led to decreased apoptotic priming and decreased, but still detectable, BCL-X_L_ dependence in circulating platelets one and two days after treatment, returning to baseline (equivalent to vehicle-treated animals) on the third day. These data, combined with our detection of short-term decreases in platelet counts after DT2216 administration, suggest that newly-produced platelets that are recently released into circulation are more resistant to apoptosis. This can be due to lower or higher expression of pro-apoptotic or pro-survival proteins, respectively. We also detected dependence on MCL-1 in these newly-formed platelets, which was not detected in vehicle-treated animals. Taken together, these data suggest that MCL-1 expression is higher in newly-formed platelets, which protects them from BCL-X_L_ targeted therapies to some extent.

Given that megakaryocytes in the bone marrow express both BCL-X_L_ and MCL-1^56^, the mitochondria-containing platelets they shed into circulation likely contain high expression levels of both these proteins. However, due to the short half-life of MCL-1^57^, this protein is expected to be rapidly lost in circulating platelets, resulting in sole dependence on BCL-XL. Thus, the ability of DT2216 therapy to avoid chronic thrombocytopenia is predicated upon both the lower VHL expression of platelets and also the dosing schedule of this agent, which maintains degradation and thus inhibition of BCL-X_L_ in tumor cells with just twice weekly administration while allowing apoptosis-resistant, MCL-1 expressing, newly formed platelets to be released into circulation. These findings also help explain why lead-in dosing with ABT-263 is able to reduce the severity of thrombocytopenia to some extent in patients, but not enough to avoid thrombocytopenia that can limit dosing^52^. Based on our data, we expect DT2216 treated patients to potentially experience transient thrombocytopenia that is then stabilized after new platelet production is enhanced. Interestingly, thrombopoietin agonists such as Eltrombopag or romiplostim that stimulate production of new, apoptosis-resistant platelets may help prevent even transient thrombocytopenia in DT2216 treated patients.

Our studies demonstrate the existence of endogenous and therapy-enhanced dependence of ovarian cancers on the pro-survival protein BCL-X_L_ and provide critical guidance for targeting this protein safely and effectively using the DT2216 PROTAC. We show, for the first time, that HGSOC cells can adapt to treatment with BH3 mimetics targeting BCL-X_L_, resulting in higher expression of resistance-promoting BCL-2 or MCL-1. This is consistent with the selection for overexpression of untargeted pro-survival proteins reported with BH3 mimetics in other cancer types^58,59^. Importantly, this adaptation also involved upregulation of BCL-X_L_ in response to ABT-263 but not DT2216 – this demonstrates that proteasome-mediated degradation of BCL-X_L_ with DT2216 may reduce the likelihood of therapy resistance arising compared with ABT-263. It is important to note that across multiple experimental models, DT2216 proved to be more effective at inducing apoptosis than ABT-263. Nevertheless, the apoptotic adaptation in HGSOC suggests that single-agent treatment with BCL-X_L_ targeting agents including DT2216 is unlikely to lead to durable responses in patients with HGSOC and likely other solid cancer types. We found that paclitaxel helped prevent apoptotic adaptation via several different mechanisms, each leading to a sensitization to DT2216. First, paclitaxel treatment led to decreased MCL-1 levels, as was evident in the OVCAR3 and OVCAR4 cell lines as well as tumors and DF83 PDX model. This was especially evident in cells that are arrested in mitosis – these cells exhibit a complete absence of MCL-1 expression and are particularly vulnerable to BCL-X_L_ inhibition. This decrease in MCL-1 within mitotically arrested cells likely forestalled the development of resistance to DT2216, leading to the enhanced responses we observed, and is consistent with previous reports showing MCL-1 phosphorylation and degradation during prolonged mitotic arrest^60,61^. Second, paclitaxel treatment also resulted in phosphorylation of BCL-X_L_ across most HGSOC cells, as evidenced by the appearance of the higher molecular weight band in the western blots and confirmed with the phosphorylation-specific antibody. This phosphorylation of BCL-X_L_ at Ser-62 has been previously reported to prevent BCL-X_L_ binding to BAX^62^, which would result in a further reduction of anti-apoptotic activity of total BCL-X_L_. There was no observed difference in DT2216-induced degradation of phosphorylated versus non-phosphorylated BCL-X_L_. Our immunoprecipitation analysis confirmed that paclitaxel and DT2216 treatment frees pro-apoptotic proteins BIM, BAX and BAK from being bound by BCL-X_L_, enabling rapid and complete commitment to apoptotic cell death. In vitro, these effects led to synergistic induction of apoptosis in HGSOC cells. In vivo, these effects enhanced tumor control in the combination treated mice compared with single agents, leading to the eradication of xenograft tumors in the recurrent HGSOC DF83 PDX model. Our results provide rationale for the evaluation of DT2216 in combination with paclitaxel for patients with recurrent HGSOC in clinical trials. These trials may benefit from the use of BH3 profiling to predict patient responses and personalize therapy since we demonstrated that BCL-X_L_ dependence can be reliably detected with this assay.

We have previously reported that HGSOC tumors are sensitive to pro-apoptotic stimuli^17^, helping to set the stage for successful deployment of DT2216 in combination with paclitaxel in this disease. It is important to note that the near-ubiquitous nature of *TP53* mutations in HGSOC impair the chemo- and radio-sensitivity of these cancers but because BH3 mimetics engage the apoptosis pathway downstream of p53 signaling, these agents can maintain high levels of efficacy despite lack of p53 signaling. Indeed, we found that a diverse array of primary tumors, PDX and PDO models were sensitive to BCL-X_L_ inhibition, suggesting that the activity of this agent will be broad, including treatment-naïve as well as heavily pretreated patients. Also, since many of the targeted anti-cancer agents currently in development will also produce pro-apoptotic signaling in ovarian cancer cells, we expect BH3 mimetics targeting BCL-X_L_ to have an enduring impact on the clinical care of these patients. Finally, although our studies focused on HGSOC, we expect DT2216 to be highly effective in other subtypes of ovarian cancer and the many other solid cancer types that have been reported to exhibit BCL-X_L_ dependence including cancers of the lung^63^, pancreas^64^, colon^65^, breast^66^, kidney^67^ and others^68^.

## Materials and Methods

### Cell and organoid culture

OVCAR4, KURAMOCHI, OVSAHO OVKATE, and JHOM-1 ovarian cancer cell lines were cultured in RPMI medium (Thermo Fisher Scientific) supplemented with 10% fetal bovine serum (FBS; Life Technologies) and 1% penicillin-streptomycin (Life Technologies). OVCAR3 cells were maintained in RPMI medium supplemented with 20% FBS and 1% penicillin-streptomycin. OV-90, OAW28, OVCAR8 and TYK-nu ovarian cancer cell lines were cultured in DMEM medium (Thermo Fisher Scientific) supplemented with 10% fetal bovine serum and 1% penicillin-streptomycin. High-grade serous ovarian cancer (HGSOC) primary tumors and patient-derived xenograft models were grown in RPMI supplemented with 10% FBS and 1% penicillin-streptomycin. The patient-derived organoids (PDO) were maintained as previously described^69^. Human and mouse peripheral blood mononuclear cells (PBMCs) were maintained in RPMI medium with 10% FBS and 1% penicillin-streptomycin. Human and mouse platelets were kept in Hank’s Balanced Salt Solution (HBSS) on a rocker at room temperature. All research involving patient samples was conducted in accordance with a Dana-Farber/Harvard Cancer Center Institutional Review Board-approved protocol, with written informed consent obtained from all participants.

### BH3 profiling

BH3 profiling was performed using flow cytometry according to established protocols^33,70^. Briefly, cultured cells were treated with trypsin (Gibco), collected and centrifuged at 200 g for 5 minutes. The cell pellet was resuspended in Mannitol Experimental Buffer 2 (MEB2), which consists of 10 mM HEPES (pH 7.5), 150 mM mannitol, 150 mM KCl, 1 mM EGTA, 1 mM EDTA, 0.1% BSA, and 5 mM succinate. The resuspended cells were subsequently added to wells of prepared plates containing the indicated peptide conditions and 0.001% digitonin. Alamethicin (25 μM) and DFNA5 (10 μM) were used as positive controls, while PUMA2A (10 μM) and DMSO were used as negative controls. After incubation for 60 minutes at 28°C, cells were fixed with a 1:4 dilution of 8% paraformaldehyde (2% final) for 15 minutes, and then neutralized with N2 buffer (1.7 M Tris base, 1.25 M glycine, pH 9.1). Cells were stained overnight with DAPI (1:1000, Abcam) and anti-Cytochrome c-Alexa Fluor 647 (1:2000, clone 6H2.B4, Biolegend) in a Tween20-based intracellular stain buffer (0.2% Tween20, 1% BSA). Cytochrome c release in response to BH3 peptide treatment was measured using an Attune NxT flow cytometer equipped with an autosampler (Thermo Fisher Scientific).

### In vitro drug treatments and Annexin V (AxV) / Propidium iodide (PI) viability assay

Cells were initially plated at a density of 10^4^ cells per well in 100 μL of culture media in flat-bottom 96-well plates (Denville). Following plating, cells were treated with various drugs at the specified concentrations: ABT-199 (1, 0.1, 0.01 μM), ABT-263 (1, 0.1, 0.01 μM), A-1331852 (1, 0.1, 0.01 μM), S63845 (1, 0.1, 0.01 μM), DT2216 (1, 0.1, 0.01 μM), ABT-737 (1, 0.1, 0.01 μM), Carboplatin (25, 100, 250 μM), Paclitaxel (0.3, 1, 3, 10, 30, 100, 300 nM), and QVD-OPH (10 μM). Combination treatments were administered simultaneously. DMSO (1%) was used as negative control. After 24, 48 or 72 hours of treatment under standard tissue culture conditions, the medium was collected and transferred to a fresh 96-well V-bottom plate (Sigma Aldrich). 25 μL of 0.0025% Trypsin (Gibco) was added to each well on the original plate and incubated for 5 minutes, trypsinized cells were then transferred to the V-bottom plate containing the collected medium. Cell staining was performed using viability markers Annexin V (AxV) and propidium iodide (PI) according to the following protocol. A staining solution was prepared using 10x Annexin Binding Buffer (0.1 M HEPES, 1.4 M NaCl, 25 mM CaCl_2_, pH 7.4, sterile filtered) mixed with AlexaFluor488-conjugated AxV and PI (Abcam) at a dilution of 1:500. The staining solution was added to the cells at a final dilution of 1:10 and incubated for 20 minutes on ice in the dark. AxV/PI positivity was then measured using an Attune NxT flow cytometer.

### Organoid CTG synergy assay

A total of 10,000 cells were seeded per well in a 96-well plate within a 10 μL Matrigel dome and treated with DT2216 and paclitaxel (PTX) for one week. The media containing the drugs was refreshed every other day. PTX concentrations ranged from 0 to 30 nM (0, 1.5, 3, 10, 30 nM), while DT2216 concentrations ranged from 0 to 1000 nM (0, 10, 30, 100, 300, 1000 nM). Synergy between the two drugs was evaluated using the SynergyFinder web application (https://synergyfinder.fimm.fi/) with calculations based on the Zero-interaction model, where a synergy scores above 10 indicates a synergistic interaction. For the apoptosis assay, 45,000 cells were seeded per well in a 48-well plate in a 15 μL Matrigel dome and similarly incubated with PTX at 1.5 nM and DT2216 at 10 nM for one week, with media changes every other day. Apoptosis was then assessed using FITC-Annexin V and 7-AAD by flow cytometry.

### Western blot analysis

Cells were seeded in 6-well plates and treated with drugs when they reached approximately 70% confluency. Cells were harvested by washing twice with cold PBS and lysed in Radio Immuno Precipitation Assay (RIPA) buffer (New England Biolabs) containing protease and phosphatase inhibitor (Thermo Scientific, 78446). Lysates were centrifuged, and supernatants were collected. Protein concentration was determined using loading was measured by Pierce™ Microplate BCA Protein Assay Kit (Thermo Fisher Scientific). Protein samples (5 μg) were resolved on 4% to 12% SDS-PAGE gels and subsequently transferred to PVDF membranes. Membranes were blocked in 5% milk and incubated with primary antibodies at 4°C overnight. After washing with PBST, membranes were incubated with the appropriate secondary antibodies for 2 hours at room temperature. Following further washes, membranes were incubated with SuperSignal West Femto Stable Peroxide & Luminol/Enhancer (Thermo Fisher Scientific) and developed using the ECL Detection System (Thermo Fisher Scientific). The following antibodies were used: BCL-2 (CST, #3498, RRID: AB_1903907, 1:500), BCL-XL (CST, #2764, RRID: AB_2228008, 1:1000), Phospho-BCL-XL (S62) (Invitrogen, #44-428G, RRID: AB_2533650, 1:1000), MCL-1 (CST, #94296, RRID: AB_2722740, 1:1000), PUMA (CST, #98672S, RRID: AB_3096180, 1:1000), BIM (CTS, #2933, RRID: AB_1030947, 1:1000), BAX (CST, #2772, RRID: AB_10695870, 1:1000), BAK NT (Millipore 06-536, RRID: AB_310159, 1:1000), BIM (CST, #2933, RRID: AB_1030947, 1:1000); BID (CST, #2002, RRID: AB_10692485, 1:1000), BAD (ROCKLAND, #200-401-Z19, RRID: AB_2612247, 1:1000), BCL-w (CST, #2724S, RRID: AB_10691557, 1:1000), Noxa (CST, #14766S, RRID: AB_2798602, 1:1000), Phospho-Histone H3 (Ser10) (CST, #9701, RRID: AB_331535, 1:1000), GAPDH (CST, #5174, RRID: AB_10622025). Anti-rabbit IgG (H1L) HRP conjugate (Thermo, 31460, RRID: AB_228341, 1:4000), and Anti-mouse IgG (H1L) HRP conjugate (Promega, W4021, RRID: AB_430834, 1:7000).

### Time lapse microscopy

OVCAR4 cells were seeded in a 96-well flat plate (CORNING, 3603) at a density of 3,000 cells per well. The following day, cells were stained with Tetramethylrhodamine, Ethyl Ester, Perchlorate (TMRE, T669) at a 1:100 dilution from a 10 µM stock solution, and CellEvent™ Caspase-3/7 Green Detection Reagent at (C10433) at a 1:100 dilution. After a 30-minute incubation at 37°C in a humidified incubator with 5% CO[, cells were treated with paclitaxel at 100 nM. The plate was then transferred to the Cellcyte live-cell imaging system (Discover Echo), and time-lapse images and videos were captured using red and green fluorescence channels to visualize mitochondrial membrane potential changes and caspase-3/7 activation, respectively.

### Mitotic Shake-Off

OVCAR3 cells were plated in a T175 cm² flask at a density of 5.0 × 10[ cells per flask. The following day, the cells were treated with paclitaxel (20 nM) for 24 hours to induce mitotic arrest. After the incubation, the medium was gently aspirated, and the cells were washed once with PBS to remove residual paclitaxel and serum. Fresh RPMI medium was added, and the flask was placed on a shaker set to 300 rpm for 5 minutes to collect loosely attached cells (shake-off). Detached mitotic cells were collected into a clean centrifuge tube. The attached cells were then detached by adding trypsin. After neutralizing the trypsin with RPMI medium containing serum, the non-mitotic cells were collected. Both mitotic and non-mitotic cells were counted using a hemocytometer or an automated cell counter. The mitotic and non-mitotic fractions were then used for BH3 profiling and Western blot analysis as required.

### Protein Immunoprecipitation

Cells were seeded in 6-well plates and treated with drugs when they reached approximately 70% confluency. After treatment, the cells were harvested by washing twice with cold PBS and then lysed on ice using in Tris Lysis Buffer (R60TX-3) containing a Protease Inhibitor Cocktail (Meso Scale Diagnostics, R70AA-1). The lysates were centrifuged, and the supernatants were collected. Protein concentration was determined using the Pierce™ Microplate BCA Protein Assay Kit (Thermo Fisher Scientific). Based on the protein concentration, 200-600 μg of protein was aliquoted and adjusted to a final volume of 250 μL with Tris Lysis Buffer for “Input” group. For immunoprecipitation, 200-600 μg of protein was combined with Tris Lysis Buffer to a total volume of 120 μL, followed by the addition of 50 μL of diluted antibody and 50 μL of Pierce Protein A/G Magnetic Beads (Thermo Fisher). Mouse anti-human BCL-X_L_ antibodies (EMD Millipore MAB3121) were used for BCL-X_L_ immunoprecipitation, and Mouse anti-human MCL-1 antibodies (BD Pharmingen 559027) were used for MCL-1 immunoprecipitation. The mixtures were incubated overnight on a rocker in the cold room to allow binding. Following incubation, the immunoprecipitated complexes were separated using magnetic separation, and the beads were washed three times with Tris Lysis Buffer on ice. Proteins were eluted by heating the samples at 95°C for 10 minutes in Tris Lysis Buffer with LDS Sample Buffer 4X (Invitrogen). A representative aliquot of the normalized whole-cell lysate was saved for Western blot analysis.

### Mouse xenograft studies

Animal experiments were performed using NSG mice (Jackson Laboratories) in accordance with the policies and regulations established by the Institutional Animal Care and Use Committee (IACUC) of the Harvard TH Chan School of Public Health, under protocol #IS00001059. For the xenograft models, cell suspensions of OVCAR3 or DF-83 were prepared in a 1:1 mixture of Matrigel and PBS. A total of 1 × 10^6^ cells were injected unilaterally into the subcutaneous space of the flanks and allowed to grow to approximately 65-100 mm^3^. Mice were then randomized into different treatment groups. The Paclitaxel cohort received intraperitoneal injection of 15 mg/kg every four days for four doses. The DT2216 cohort received intraperitoneal injection of 15 mg/kg every four days. The combination cohort received both Paclitaxel and DT2216, while the vehicle cohort received corresponding vehicles. Tumor growth and body weight were monitored every other day using electronic calipers, and tumor volume was calculated using the formula V = 4/3 × π × Length/2 × (Width/2)^2^. Blood samples were collected via tail vein every four days for OVCAR3 xenograft models and every seven days for DF-83 xenograft models, and blood cell counts were performed. Mice were euthanized when tumors reached the predetermined endpoint. Tumors and healthy tissues were collected for further analysis. Healthy tissues were rinsed with Hank’s Balanced Salt Solution and placed into 4% formaldehyde solution (Electron Microscopy Sciences) for 24 hours at 4°C. Sections were then stained with hematoxylin and eosin stain, and microphotographs of the sections were captured using a Leica DM2000 microscope equipped with a 40X objective.

### BH3 Profiling of Primary Patient Samples and Xenograft tissues

Tissue samples were initially minced into approximately 1mm^3^ pieces using scalpels and then subjected to enzymatic dissociation with collagenase/dispase for 30 minutes at 37°C. The resulting cell suspensions were then filtered through a 100 mm filter to obtain single-cell preparations, which were then centrifuged at 200g for 5 minutes at 4°C to pellet the cells. The cell pellet was resuspended and cultured in RPMI1640 supplemented with 10% FBS and 1% penicillin-streptomycin for 24 hours in the absence or presence of the specified drugs. Following culture, cells were collected and stained with specific antibodies: anti-human EpCAM for primary patient and xenograft samples, anti-human or anti-mouse CD61 for platelets, and anti-human or anti-mouse CD45 for PBMCs. Subsequently, Flow cytometry based BH3 profiling was then performed. PBMCs were isolated using anti-human CD45 staining in primary patient and anti-mouse CD45 for xenograft samples, and the remaining cell identity stains were analyzed on CD45 negative-gated cells.

### Isolation of PBMCs and platelets

Peripheral blood was collected from human donors in compliance with an IRB-approved protocol at Harvard University, using lavender-top tubes containing EDTA. The peripheral blood was diluted with an equal volume phosphate-buffered saline (PBS) and then carefully layered over Ficoll-Paque (GE Healthcare). After centrifugation at 177 g for 20 minutes at room temperature with low brake settings, the supernatant enriched with platelets was transferred to a new tube. Prostaglandin E1 (PGE1) was added to achieve a final dilution of 1:10,000 from a 10mM stock solution. The solution was then centrifuged at 1000 g for 5 minutes at room temperature. The resulting pellet, which contained platelets, was resuspended with HBSS and counted. The interface layer, containing PBMCs interface, was carefully aspirated and washed with PBS before counting. The isolated PBMCs and platelets were subsequently used for BH3 profiling and chemosensitivity assay.

### Colony formation assay

OVCAR3 cells were trypsinized and resuspended in RPMI1640 supplemented with 20% FBS and 1% penicillin-streptomycin. Following cell counting, 600 cells per well were seeded into 24-well plates. Cells were then treated with the indicated conditions the following day. After 24 hours of exposure, the medium was replaced with fresh medium, and the cells were incubated for additional two weeks under standard culture conditions. At the end of the incubation period, the medium was removed, and the cells were washed with PBS. The cells were subsequently stained with 0.05% crystal violet solution for 30[minutes. Following staining, the crystal violet solution was removed, and the cells were washed twice with PBS. Stained colonies were imaged using the ECL Detection System (Thermo Fisher Scientific, 32209).

### ProteinAtlas.org

Using the Human Protein Atlas online database^71^, we investigated BCL-X_L_ expression in different OvCa primary tumor specimens using the antibody HPA035734.

### PDX models

With the exception of DF09 and DF20, which were derived from therapy-naïve patients, the primary ovarian tumors used for the generation of the PDX models were isolated from women with advanced high grade serous ovarian cancer that have been treated with multiple chemotherapies. The establishment of these models, their clinical and genomic characteristics (e.g., BRCA mutations) have been previously described^45^. mCherry–Luciferase was transduced into dissociated tumor cells in vitro with a lentiviral vector and then placed under puromycin selection for 5–7 days before expansion in vivo (up to six passages) to generate viably frozen stocks. PDX models tested negative for mycoplasma contamination and were authenticated by genomic comparison with the original tumor.

### PDX ex vivo dose-response analyses

Frozen PDX ascites cells were thawed and allowed to recover for two days in a 1:1:1 ratio of Medium 199 (11150067, Life Tech), RPMI-1640 (11875119, ThermoFisher) and DMEM/F12 (11330057, ThermoFisher), supplemented with 2% HI-FBS (Corning), 1% (v/v) Penicillin-Streptomycin (Sigma-Aldrich), 1% (v/v) Insulin-Transferrin-Selenium (354352, Corning), 0.5 ng/ml of 17 beta-estradiol (US Biological), 0.2 pg/mL of triiodothyronine (Sigma-Aldrich), 0.025 ug/mL all-trans retinoic acid (Beantown Chemical), 20 ug/mL of insulin (Sigma-Aldrich), 0.5 ng/mL of EGF (Peprotech), 0.5 mg/mL hydrocortisone (Sigma-Aldrich) and 25 ng/mL of Cholera Toxin (Calbiochem). Cells were then seeded into 384-well white plates (3570, Corning) in a 1:1 ratio of MCDB105 (117-500, Westnet), M199 medium (11150067, Life Tech) supplemented with 2% HI-FBS and 1% (v/v) Penicillin-Streptomycin. After 4 hours of recovery in the incubator, cells in 384 well-plates were treated with indicated dose ranges of DT2216 and paclitaxel using a HP D300 drug printer. Dose combination studies were performed with a dose of DT2216 starting from 10 μM followed by log10 based serial dilution for nine dose ranges (0.05 - 10 μM), and a same log10 based serial dilution for paclitaxel (0.05 - 10 μM) for five dose ranges. DMSO was used as vehicle control. After 96 hours of drug treatment, cell viability was measured by luciferase assay with a PerkinElmer Envision plate reader (384-L1 aperture). Briefly, 20 μL of 1mg/mL D-Luciferin (LUCK-1G, GoldBio) in DPBS was added to each well of the 384-well plates, and then plates were shaken for 1 minute and incubated for 1 minute before reading. To determine growth rate, a parallel plate was read on the day of drug addition to calculate the luciferase signal before treatment (Day 0).

### Statistical analysis

Significance was tested by student’s t test, One-Way or Two-Way ANOVA with Holm-Sidak’s correction for multiple hypothesis testing using GraphPad PRISM software.

## Data availability

The raw data generated in this study are available upon request from the corresponding author.

## Acknowledgments

We thank the members of our labs for their thoughtful suggestions on this work.

## Funding

This work was supported by funding from:

National Institutes of Health grant R00CA188679 (KAS)

National Institutes of Health grant R37CA248565 (KAS)

National Institutes of Health grant R01DK125263 (KAS)

National Institutes of Health grant F31CA246811 (SY)

National Institutes of Health grant 1F31CA281083 (LJ)

National Institutes of Health grant F31CA275321 (MHF)

Alex’s Lemonade Stand Foundation (KAS)

Andrew McDonough B+ Foundation (KAS)

Blavatnik Institute at Harvard (KAS)

HSPH Dean’s Fund for Scientific Advancement (KAS)

HSPH National Institute for Environmental Health Sciences (NIEHS) Center Grant National Institutes of Health grant P30ES000002 (KAS)

National Institutes of Health grant K08CA237871 (EHS)

Ovarian Cancer Research Alliance (EHS and KAS)

## Author contributions

Conceptualization: XQ, KAS

Methodology: XQ, AP, LJ, YM, BSS, WX, JC, CF, FG, JS, SY, MHCF, FP, YY Investigation: XQ, AP, LJ, YM, BSS, WX, JC, CF, FG, JS, SY, MHCF, FP, YY

Visualization: XQ, KS

Funding acquisition: KAS

Project administration: KAS

Supervision: KAS

Writing – original draft: XQ, KAS

Writing – review & editing: all authors

## SUPPLEMENTARY MATERIALS

### Supplementary Figures

Figure S1 (Related to Figure 1)

Figure S2 (Related to Figure 2)

Figure S3 (Related to Figure 3)

Figure S5 (Related to Figure 5)

Figure S7 (Related to Figure 7)

## References

1. Schmid, B. C. & Oehler, M. K. New perspectives in ovarian cancer treatment. Maturitas 77, 128–36 (2014).

2. Wang, Z. C. et al. Profiles of Genomic Instability in High-Grade Serous Ovarian Cancer Predict Treatment Outcome. Clinical cancer researchJ: an official journal of the American Association for Cancer Research 18, 5806–5815 (2012).

3. Verhaak, R. G. W. et al. Prognostically relevant gene signatures of high-grade serous ovarian carcinoma. 123, 517–525 (2013).

4. Cancer Genome Atlas Research Network, T., et al. Integrated genomic analyses of ovarian carcinoma. Nature 474, 609–15 (2011).

5. Stickles, X. B. et al. In vitro analysis of ovarian cancer response to cisplatin, carboplatin and paclitaxel identifies common pathways that are also associated with overall patient survival. British Journal of Cancer 106, 1967–1975 (2012).

6. Agarwal, R. & Kaye, S. B. Ovarian cancer: strategies for overcoming resistance to chemotherapy. Nature reviews. Cancer 3, 502–16 (2003).

7. Colombo, P.-E. et al. Sensitivity and resistance to treatment in the primary management of epithelial ovarian cancer. Critical reviews in oncology/hematology 89, 207–16 (2014).

8. Fulda, S. & Debatin, K.-M. Extrinsic versus intrinsic apoptosis pathways in anticancer chemotherapy. Oncogene 25, 4798–811 (2006).

9. Johnstone, R. W., Ruefli, A. A., Lowe, S. W. & Victoria, E. M. ApoptosislJ: A Link between Cancer Genetics and Chemotherapy Defects in apoptosis underpin both tumorigenesis and. 108, 153–164 (2002).

10. Sarosiek, K. A., Ni Chonghaile, T. & Letai, A. Mitochondria: gatekeepers of response to chemotherapy. Trends in Cell Biology 1–8 (2013) doi:10.1016/j.tcb.2013.08.003.

11. Campbell, K. J. & Tait, S. W. G. Targeting BCL-2 regulated apoptosis in cancer. 1–11 (2018).

12. Meier, P., Finch, a & Evan, G. Apoptosis in development. Nature 407, 796–801 (2000).

13. Elmore, S. Apoptosis: a review of programmed cell death. Toxicologic pathology 35, 495–516 (2007).

14. Lopez, J. & Tait, S. W. G. Mitochondrial apoptosis: Killing cancer using the enemy within. British Journal of Cancer 112, 957– 962 (2015).

15. Brunelle, J. K. & Letai, A. Control of mitochondrial apoptosis by the Bcl-2 family. Journal of cell science 122, 437–441 (2009).

16. Green, D. R. & Walczak, H. Apoptosis therapylJ: driving cancers down the road to ruin. Nature Medicine 19, 131–133 (2013).

17. Ni Chonghaile, T., et al. Pretreatment mitochondrial priming correlates with clinical response to cytotoxic chemotherapy. Science 334, 1129–1133 (2011).

18. Sarosiek, K. A. et al. BID preferentially activates BAK while BIM preferentially activates BAX, affecting chemotherapy response. Molecular cell 51, 751–765 (2013).

19. Montero, J. et al. Drug-Induced Death Signaling Strategy Rapidly Predicts Cancer Response to Chemotherapy. Cell 160, 977– 989 (2015).

20. Abed, M. N., Abdullah, M. I. & Richardson, A. Antagonism of Bcl-XL is necessary for synergy between carboplatin and BH3 mimetics in ovarian cancer cells. Journal of Ovarian Research 9, 25 (2016).

21. Peale, F. V. et al. Navitoclax (ABT-263) Reduces Bcl-xL-Mediated Chemoresistance in Ovarian Cancer Models. Molecular Cancer Therapeutics 11, 1026–1035 (2012).

22. Stover, E. H. et al. Pooled genomic screens identify anti-apoptotic genes as targetable mediators of chemotherapy resistance in ovarian cancer. Molecular Cancer Research 17, 2281–2293 (2019).

23. Yokoyama, T., Kohn, E. C., Brill, E. & Lee, J. Apoptosis is augmented in high-grade serous ovarian cancer by the combined inhibition of Bcl-2 / Bcl-xL and PARP. 1064–1074 (2017) doi:10.3892/ijo.2017.3914.

24. Tai, Y. T. et al. BAX protein expression and clinical outcome in epithelial ovarian cancer. Journal of clinical oncologyJ: official journal of the American Society of Clinical Oncology 16, 2583–90 (1998).

25. Al-Alem, L. F., Baker, A. T., Pandya, U. M., Eisenhauer, E. L. & Rueda, B. R. Understanding and targeting apoptotic pathways in ovarian cancer. Cancers 11, (2019).

26. Hajra, K. M., Tan, L. & Liu, J. R. Defective apoptosis underlies chemoresistance in Ovarian Cancer. in Ovarian Cancer vol. 622 197–208 (Springer New York, New York, NY, 2002).

27. Chen, Y. et al. MicroRNA letlJ7dlJ5p rescues ovarian cancer cell apoptosis and restores chemosensitivity by regulating the p53 signaling pathway via HMGA1. International Journal of Oncology (2019) doi:10.3892/ijo.2019.4731.

28. Merino, D. et al. BH3-Mimetic Drugs: Blazing the Trail for New Cancer Medicines. Cancer Cell 34, 879–891 (2018).

29. Khan, S. et al. A selective BCL-XL PROTAC degrader achieves safe and potent antitumor activity. Nature Medicine 25, 1938– 1947 (2020).

30. He, Y. et al. DT2216—a Bcl-xL-specific degrader is highly active against Bcl-xL-dependent T cell lymphomas. Journal of Hematology & Oncology 13, 95 (2020).

31. Domcke, S., Sinha, R., Levine, D. A., Sander, C. & Schultz, N. Evaluating cell lines as tumour models by comparison of genomic profiles. Nature Communications 4, 1–10 (2013).

32. Letai, A. et al. Distinct BH3 domains either sensitize or activate mitochondrial apoptosis, serving as prototype cancer therapeutics. Cancer cell 2, 183–92 (2002).

33. Fraser, C., Ryan, J. & Sarosiek, K. BH3 Profiling: A Functional Assay to Measure Apoptotic Priming and Dependencies. in *Methods in Molecular Biology* vol. 1877 61–76 (2019).

34. Vo, T.-T. et al. Relative mitochondrial priming of myeloblasts and normal HSCs determines chemotherapeutic success in AML. Cell 151, 344–355 (2012).

35. Davids, M. S. et al. Mitochondrial Apoptotic Priming Is Associated with Clinical Response to the Bcl-2 Antagonist ABT-199 in Chronic Lymphocytic Leukemia. Blood 124, (2014).

36. Fraser, C. S. et al. Exploiting endogenous and therapy-induced apoptotic vulnerabilities in immunoglobulin light chain amyloidosis with BH3 mimetics. Nat Commun 13, 5789 (2022).

37. Inde, Z. et al. Age-dependent regulation of SARS-CoV-2 cell entry genes and cell death programs correlates with COVID-19 severity. Science Advances 7, 1–18 (2021).

38. Dempster, J. M. et al. Extracting Biological Insights from the Project Achilles Genome-Scale CRISPR Screens in Cancer Cell Lines. 720243 Preprint at 10.1101/720243 (2019).

39. Javellana, M. et al. Neoadjuvant Chemotherapy Induces Genomic and Transcriptomic Changes in Ovarian Cancer. Cancer Res 82, 169–176 (2022).

40. Joly, F. et al. A phase II study of Navitoclax (ABT-263) as single agent in women heavily pretreated for recurrent epithelial ovarian cancer: The MONAVI - GINECO study. Gynecol Oncol 165, 30–39 (2022).

41. Liu, Q., Yin, X., Languino, L. R. & Altieri, D. C. Evaluation of drug combination effect using a Bliss independence dose-response surface model. Stat Biopharm Res 10, 112–122 (2018).

42. Gillet, J.-P., Varma, S. & Gottesman, M. M. The clinical relevance of cancer cell lines. Journal of the National Cancer Institute 105, (2013).

43. Ghandi, M. et al. Next-generation characterization of the Cancer Cell Line Encyclopedia. Nature 569, 503–508 (2019).

44. Hill, S. J. et al. Prediction of DNA repair inhibitor response in short-term patient-derived ovarian cancer organoids. Cancer discovery 8, 1404–1421 (2018).

45. Liu J.F. et al. Establishment of patient-derived tumor xenograft models of epithelial ovarian cancer for preclinical evaluation of novel therapeutics. Clinical Cancer Research 23, 1263–1273 (2017).

46. Shoemaker, A. R. et al. The Bcl-2 Family Inhibitor ABT-263 Shows Significant but Reversible Thrombocytopenia in Mice. Blood 108, 1107 (2006).

47. Mitra, A. K. et al. In vivo tumor growth of high-grade serous ovarian cancer cell lines. Gynecologic oncology 138, 372–7 (2015).

48. Gandhi, L. et al. Phase I study of navitoclax (ABT-263), a novel bcl-2 family inhibitor, in patients with small-cell lung cancer and other solid tumors. Journal of Clinical Oncology 29, 909–916 (2011).

49. Johnstone, R. W., Ruefli, A. A. & Lowe, S. W. Apoptosis: A link between cancer genetics and chemotherapy. Cell 108, 153–164 (2002).

50. Sarosiek, K. A. et al. Developmental Regulation of Mitochondrial Apoptosis by c-Myc Governs Age- and Tissue-Specific Sensitivity to Cancer Therapeutics. Cancer Cell 1–15 (2016) doi:10.1016/j.ccell.2016.11.011.

51. Spetz, J. K. E. et al. Heightened apoptotic priming of vascular cells across tissues and life span predisposes them to cancer therapy-induced toxicities. Sci Adv 8, eabn6579 (2022).

52. Roberts, A. W. et al. Ongoing phase I studies of ABT-263: Mitigating Bcl-XL induced thrombocytopenia with lead-in and continuous dosing. JCO 27, 3505–3505 (2009).

53. Leverson, J. D. et al. Exploiting selective BCL-2 family inhibitors to dissect cell survival dependencies and define improved strategies for cancer therapy. Science translational medicine 7, 279ra40 (2015).

54. Desai, P. et al. A Phase 1 First-in-Human Study of the MCL-1 Inhibitor AZD5991 in Patients with Relapsed/Refractory Hematologic Malignancies. Clinical Cancer Research 30, 4844–4855 (2024).

55. Zervantonakis, I. K. et al. Systems analysis of apoptotic priming in ovarian cancer identifies vulnerabilities and predictors of drug response. Nature Communications 8, 365 (2017).

56. Debrincat, M. et al. Mcl-1 and Bcl-x(L) coordinately regulate megakaryocyte survival. Blood 119, 5850–8 (2012).

57. Opferman, J. T. Unraveling MCL-1 degradation. Cell Death & Differentiation 13, (2006).

58. Choudhary, G. S. et al. MCL-1 and BCL-xL-dependent resistance to the BCL-2 inhibitor ABT-199 can be overcome by preventing PI3K/AKT/mTOR activation in lymphoid malignancies. Cell Death and Disease 6, e1593 (2015).

59. Ramsey, H. E. et al. A Novel MCL-1 Inhibitor Combined with Venetoclax Rescues Venetoclax Resistant Acute Myelogenous Leukemia. Cancer Discovery CD-18–0140 (2018) doi:10.1158/2159-8290.CD-18-0140.

60. Chu, R., Terrano, D. T. & Chambers, T. C. Cdk1/cyclin B plays a key role in mitotic arrest-induced apoptosis by phosphorylation of Mcl-1, promoting its degradation and freeing Bak from sequestration. Biochem Pharmacol 83, 199–206 (2012).

61. Wertz, I. E. et al. Sensitivity to antitubulin chemotherapeutics is regulated by MCL1 and FBW7. Nature 471, 110–114 (2011).

62. Upreti, M. et al. Identification of the Major Phosphorylation Site in Bcl-xL Induced by Microtubule Inhibitors and Analysis of Its Functional Significance. J Biol Chem 283, 35517–35525 (2008).

63. Park, D. et al. Novel small-molecule inhibitors of Bcl-XL to treat lung cancer. Cancer Research 73, 5485–5496 (2013).

64. Ikezawa, K. et al. Increased Bcl-xL Expression in Pancreatic Neoplasia Promotes Carcinogenesis by Inhibiting Senescence and Apoptosis. Cell Mol Gastroenterol Hepatol 4, 185–200.e1 (2017).

65. Dunne, P. D. et al. Bcl-xL as a poor prognostic biomarker and predictor of response to adjuvant chemotherapy specifically in BRAF-mutant stage II and III colon cancer. Oncotarget 9, 13834–13847 (2018).

66. Alcon, C. et al. ER+ Breast Cancer Strongly Depends on MCL-1 and BCL-xL Anti-Apoptotic Proteins. Cells 10, 1659 (2021).

67. Grubb, T. et al. A Mesenchymal Tumor Cell State Confers Increased Dependency on the BCL-XL Antiapoptotic Protein in Kidney Cancer. Clin Cancer Res 28, 4689–4701 (2022).

68. Diepstraten, S. T. et al. The manipulation of apoptosis for cancer therapy using BH3-mimetic drugs. Nature Reviews Cancer 22, 45–64 (2022).

69. Matoba, Y. et al. Targeting Galectin 3 illuminates its contributions to the pathology of uterine serous carcinoma. Br J Cancer 130, 1463–1476 (2024).

70. Ryan, J., Montero, J., Rocco, J. & Letai, A. iBH3: simple, fixable BH3 profiling to determine apoptotic priming in primary tissue by flow cytometry. Biological Chemistry 0, (2016).

71. Uhlén, M. et al. A human protein atlas for normal and cancer tissues based on antibody proteomics. Molecular & cellular proteomicsJ: MCP 4, 1920–32 (2005).

