## Supplemental Figures and Tables for "Paclitaxel-induced mitotic arrest results in a convergence of apoptotic dependencies that can be safely exploited by BCL-X_L_ degradation to overcome cancer chemoresistance"

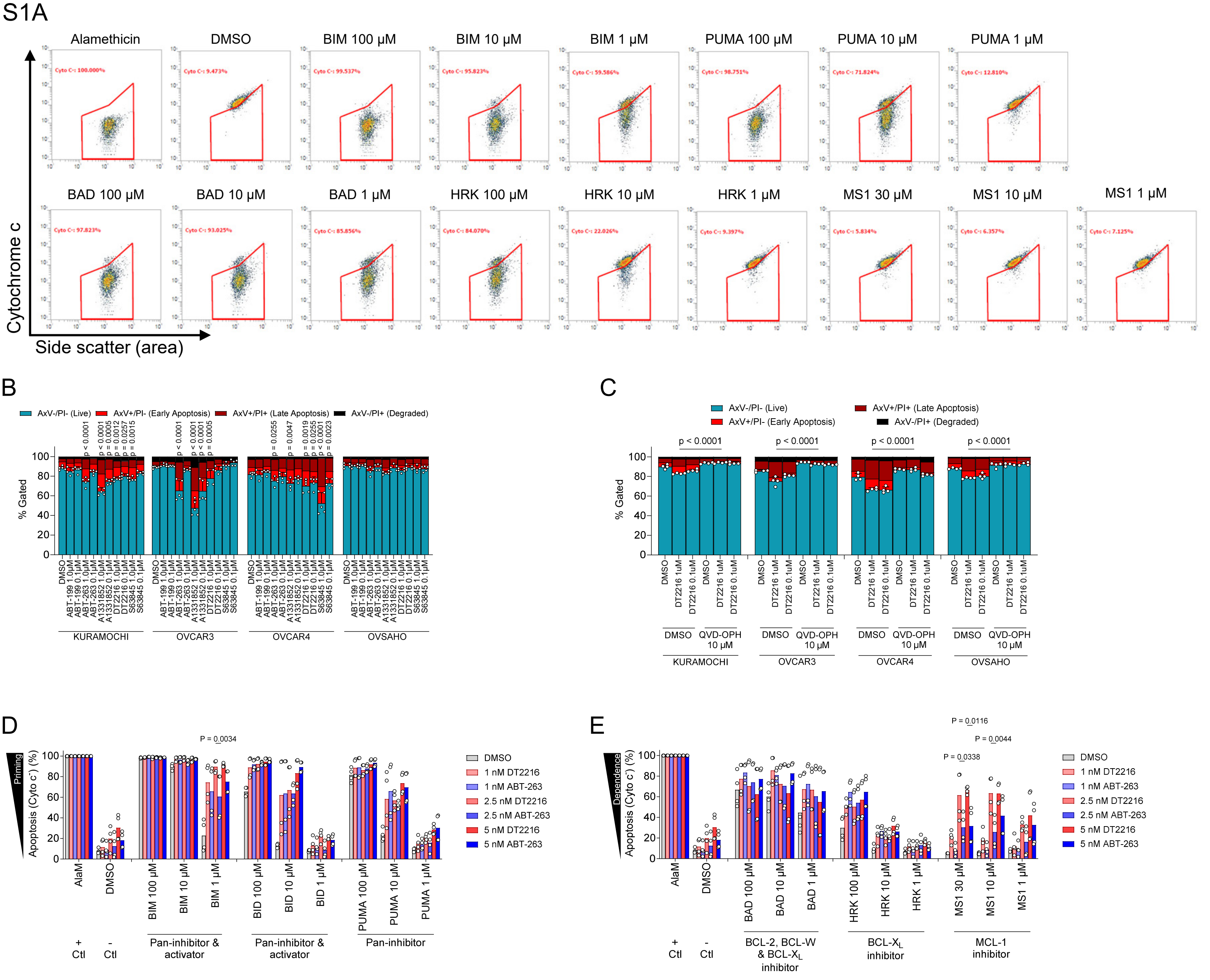


**Figure S1 (Related to Figure 1): Caspase inhibition prevents apoptosis execution in response to DT2216 treatment.** (A) Representative flow cytometry dot plots assessing percentage of cells that have released cytochrome c in response to pro-apoptotic BH3 peptides. (Representative data of *n* = 3 biological replicates). (B) Annexin V/PI staining and flow cytometry analysis of OvCa cell lines treated with single agent BH3 mimetics. Mean ± SEM is shown from n = 3 biological replicates. (C) Annexin V/ PI staining and flow cytometry analysis of OvCa cell lines treated with DT2216 (degrades BCL-XL) for 72 hours alone or in the presence of the pan-caspase inhibitor QVD-OPH. Mean ± SEM is shown for *n* = 4 biological replicates. (D-E) BH3 profiling of KURAMOCHI cells treated with indicated dose of DT2216 or ABT-263 for 96 hours to measure apoptotic priming (D) or dependencies (E). Mean ± SEM is shown for n = 4 biological replicates. P values were calculated using two-way ANOVA with Holm-Sidak's multiple comparisons test.

S2A

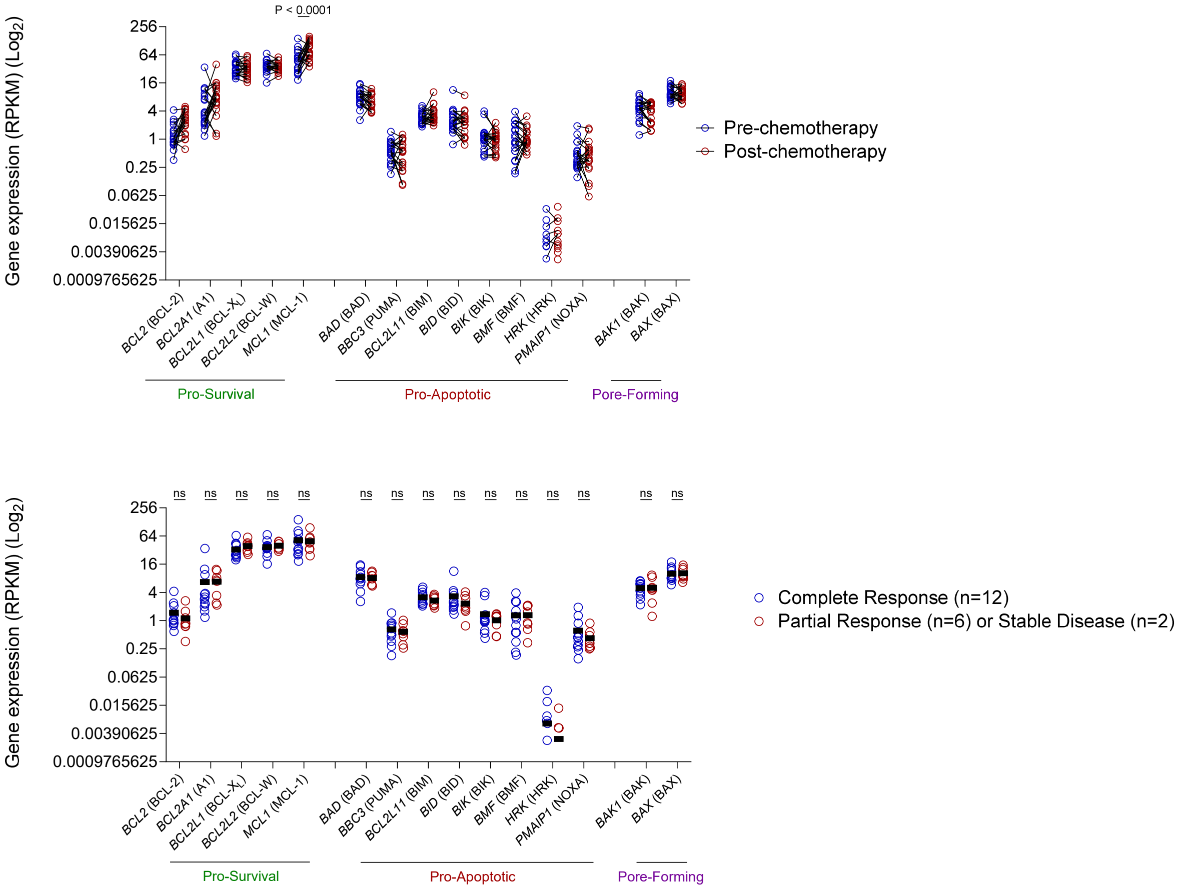


**Figure S2 (Related to Figure 2): Neoadjuvant therapy is associated with increased levels of MCL-1.** (A) Gene expression analysis from RNA-seq performed on HGSOC primary tumors before and after administration of neoadjuvant chemotherapy. (*n* = 20 tumors per group; two-way ANOVA with Holm-Sidak's multiple comparisons test). (B) Gene expression analysis from RNA-seq performed on HGSOC primary tumors before administration of neoadjuvant chemotherapy comparing patients that achieved a complete response versus those with partial response or stable disease. (*n* = 20 tumors per group; two-way ANOVA with Holm-Sidak's multiple comparisons test).

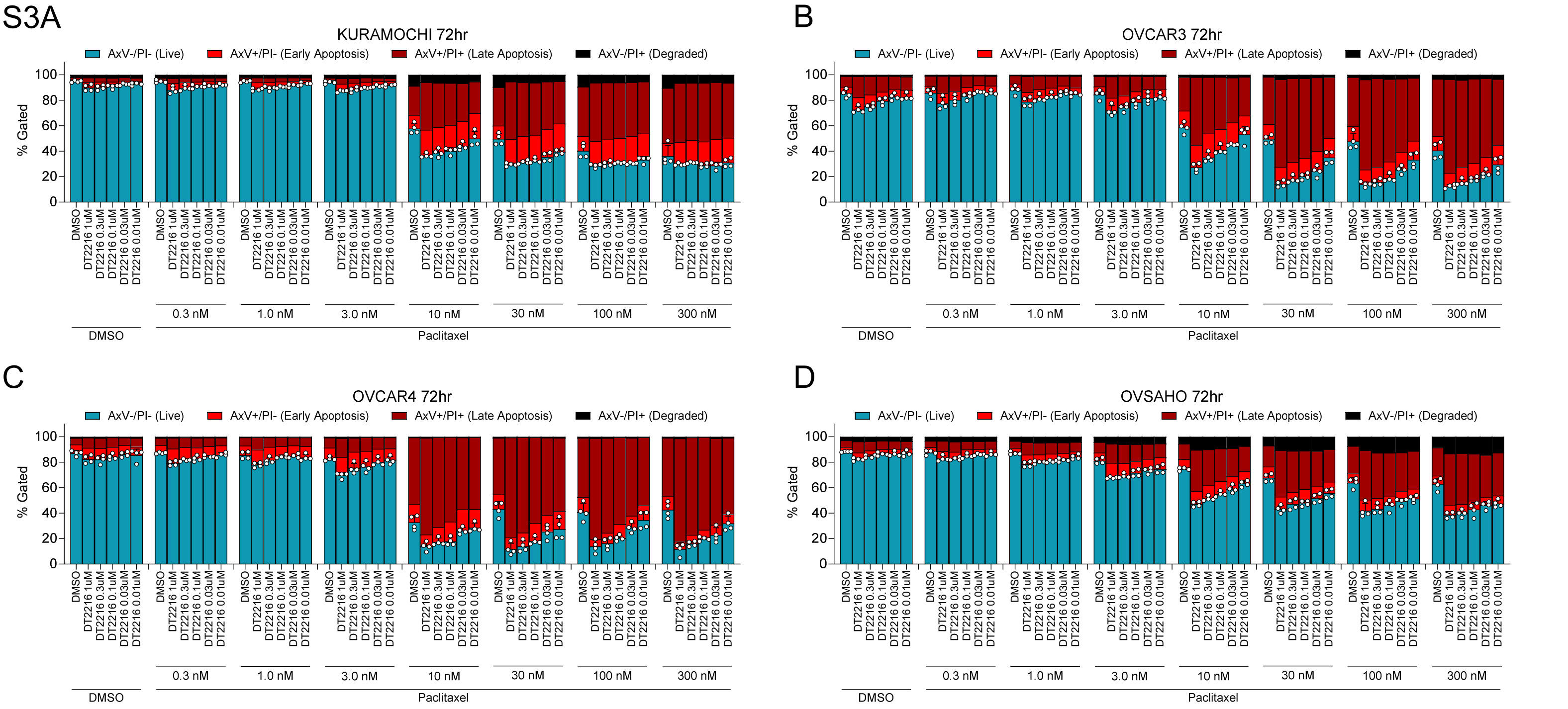


**Figure S3 (Related to Figure 3): DT2216 enhances OvCa cell line sensitivity to paclitaxel.** (A-D) Annexin V/PI staining and flow cytometry analysis of OvCa cell lines treated with agents targeting BCL-2 family proteins including ABT-199 (inhibits BCL-2 only), ABT-263 (inhibits BCL-2, BCL-XL and BCL-W), DT2216 (degrades BCL-XL), A1331852 (inhibits BCL-XL), or S63845 (inhibits MCL-1) for 72 hours alone or in combination with paclitaxel. (*n* = 4 biological replicates).

S5A

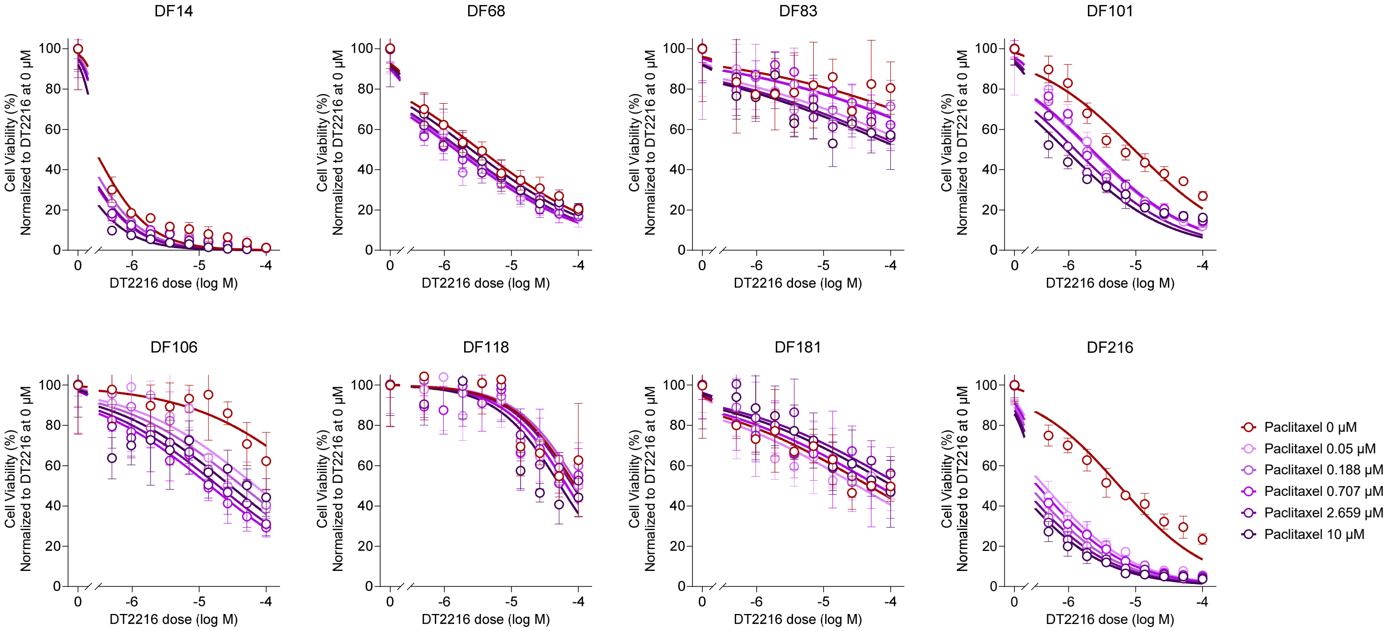


B

| | Synergy Score | | | Most synergistic area score | | |
|---|---|---|---|---|---|---|
| PDX model | Experiment 1 | Experiment 2 | Average | Experiment 1 | Experiment 2 | Average |
| DF14 | 4.753 | 8.708 | 6.7305 | 18.38 | 15.73 | 17.055 |
| DF68 | 8.621 | 13.23 | 10.9255 | 14.04 | 23.76 | 18.9 |
| DF83 | 12.499 | 13.278 | 12.8885 | 19.37 | 23.74 | 21.555 |
| DF101 | 21.557 | 6.776 | 14.1665 | 27.65 | 9.76 | 18.705 |
| DF106 | 24.821 | 7.883 | 16.352 | 38.83 | 25.36 | 32.095 |
| DF118 | 3.805 | 20.363 | 12.084 | 11.55 | 25.32 | 18.435 |
| DF181 | 3.721 | 1.218 | 2.4695 | 9.98 | 14.66 | 12.32 |
| DF216 | 30.905 | 9.838 | 20.3715 | 39.85 | 20.16 | 30.005 |

C

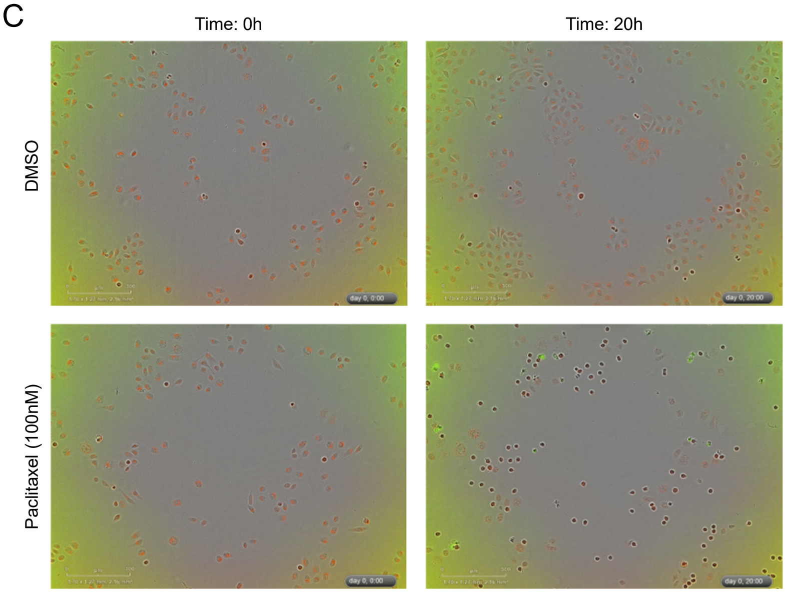


**Figure S5 (Related to Figure 5): HGSOC PDX models are broadly sensitive to combination treatment with paclitaxel and DT2216.** (A) OvCa PDX models collected during in vivo growth were cultured ex vivo for 48 hours, then treated with DT2216 ± paclitaxel and viability was measured after 96 hours by adding luciferin and measuring luminescence. Viability measurements were normalized to paclitaxel treatment only (DT2216 0 µM) to compare effects of DT2216 alone versus in combination with paclitaxel. (Representative data of $n$ = 2 biological replicates). (B) Overall synergy scores and most synergistic area scores for PDX models treated in (A) across two biological replicates. (C) Representative images from time lapse microscopy of OVCAR4 cells treated with DMSO or paclitaxel at 100nM for 20 hours.

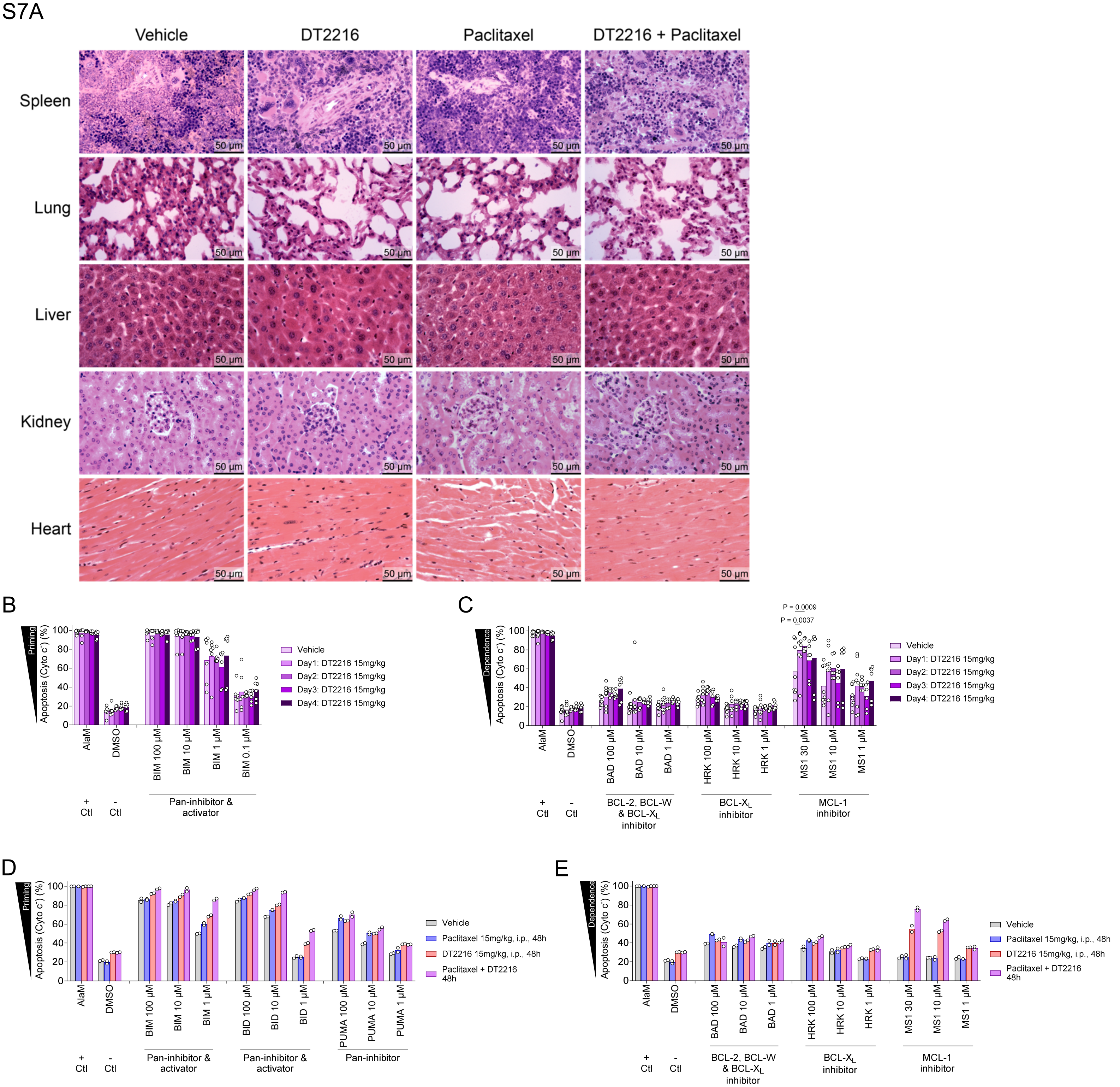


**Figure S7 (Related to Figure 7): DT2216, paclitaxel and combination treatment does not cause detectable signs of drug toxicity in healthy tissues.** (A) The spleen, lung, liver, kidney, and heart tissues show preserved structural integrity with no signs of cellular disruption or necrosis; the lung alveoli, liver hepatocytes and sinusoids, kidney glomeruli and tubules, and heart cardiomyocytes all maintain normal histological appearances. The splenic architecture is intact with defined white and red pulp, liver sections show uniform hepatocytes and open sinusoids, while cardiomyocytes display regular striations without degeneration. (Representative data for *n* = 5 independent animals). (B-C) BH3 profiling of PBMCs isolated from mice treated with indicated agents to measure apoptotic priming (B) or dependencies (C). (n = 6 biological replicates; two-way ANOVA with Holm-Sidak's multiple comparisons test). (D-E) BH3 profiling of DF83 PDX model collected after 48 hour treatment with indicated agents to measure apoptotic priming (D) or dependencies (E). (n = 2 biological replicates).

| DF code | Diagnosis | Anatomical location of specimen provided | Treatments prior to specimen collection | Days since last treatment |
|---|---|---|---|---|
| DF3968 | Metastatic high grade serous carcinoma | Left perirectal tumor | 1) Cisplatin/paclitaxel; 2) carboplatin/paclitaxel; 3) doxorubicin liposomal; 4) clinical trial drug #1; 5) clinical trial drug #2 | ~14-44 |
| DF4124 | Metastatic high grade serous carcinoma | Omentum | 1) Carboplatin/paclitaxel (neoadjuvant) | On treatment |
| DF4379 | High grade serous carcinoma | Recto-vaginal septum | 1) Carboplatin/paclitaxel | 694 |
| DF4411 | Metastatic high grade serous carcinoma | Omentum | 1) Carboplatin/paclitaxel (neoadjuvant) | On treatment |
| DF4423 | High grade mullerian carcinoma; favor high grade serous carcinoma | Omentum | None | N/A |
| DF4225 | Metastatic high grade serous carcinoma | Omentum | 1) Paclitaxel; 2) carboplatin/paclitaxel | 569 |
| DF5053 | High grade serous carcinoma | Tubo-ovarian tumor | 1) Carboplatin/paclitaxel (neoadjuvant) | On treatment |

**Table 1: Ovarian cancer primary tumors.**

| PDO | Treatment | Histology | Sites | Germline | Copy Number Alterations |
| --- | --- | --- | --- | --- | --- |
| 17-39 | Recurrent | HGSOC | Multiple | *BRCA1* | *Myc Amp* |
| 17-116 | Neoadjuvant | HGSOC | Omentum | | |
| 17-121 | Recurrent | HGSOC | Pleural Effusion | | |
| 18-47 | Untreated | HGSOC | Omentum | | *CCNE1 Amp* |
| TC1005 | Recurrent | HGSOC | | | |

**Table 2: Ovarian cancer patient-derived organoids.**

| PDX Model | Histo-logical Subtype | Source | # Prior Chemo Cycles | Prior Platinum | Germline BRCA Status | PAX8 (IHC) | WT1 (IHC) | Pan-CK (IHC) | Mutations | Human CA125 |
|---|---|---|---|---|---|---|---|---|---|---|
| DF14 | HGSOC | Ascites | 5 | Yes | Unknown | Pos | Pos | Pos | Not performed | Pos |
| DF20 | HGSOC | Ascites | 0 | No | Unknown | Pos | Pos | Pos | TP53; PTEN; PPM1D | Pos |
| DF68 | HGSOC | Ascites | 5 | Yes | BRCA1 Q563X | Pos | Pos | Pos | TP53; BRCA1, PTEN LOH | Pos |
| DF83 | HGSOC | Ascites | 4 | Yes | Unknown | Pos | Neg | Pos | TP53; CDKN2A LOH | Pos |
| DF101 | HGSOC | Pleural Fluid | 2 | Yes | BRCA1 187delAG | Pos | Pos | Pos | TP53, BRCA1, NBN, PTEN LOH | Pos |
| DF106 | HGSOC | Ascites | 1 | Yes | Unknown | Pos | Pos | Pos | TP53, CDKN2A | Pos |
| DF118 | HGSOC | Ascites | 1 | Yes | Unknown | Pos | Pos | Pos | TP53 | Pos |
| DF172 | Mixed (Serous & Endometroid) | Ascites | 2 | Yes | Unknown | Pos | Patchy | Pos | TP53; CDKN2A LOH | Pos |
| DF181 | HGSOC | Ascites | 7 | Yes | WT | Neg | Neg | Pos | TP53, BRIP1 | Neg |
| DF216 | Adeno-carcinoma | Ascites | 2 | Yes | WT | Pos | Pos | Pos | TP53 | Pos |

### Table 3: Ovarian cancer PDX models.
